# Structure and function of the human apoptotic scramblase Xkr4

**DOI:** 10.1101/2024.08.07.607004

**Authors:** Sayan Chakraborty, Zhang Feng, Sangyun Lee, Omar E. Alvarenga, Aniruddha Panda, Renato Bruni, George Khelashvili, Kallol Gupta, Alessio Accardi

## Abstract

Phosphatidylserine externalization on the surface of dying cells is a key signal for their recognition and clearance by macrophages and is mediated by members of the X-Kell related (Xkr) protein family. Defective Xkr-mediated scrambling impairs clearance, leading to inflammation. It was proposed that activation of the Xkr4 apoptotic scramblase requires caspase cleavage, followed by dimerization and ligand binding. Here, using a combination of biochemical approaches we show that purified monomeric, full-length human Xkr4 (hXkr4) scrambles lipids. CryoEM imaging shows that hXkr4 adopts a novel conformation, where three conserved acidic residues create an electronegative surface embedded in the membrane. Molecular dynamics simulations show this conformation induces membrane thinning, which could promote scrambling. Thinning is ablated or reduced in conditions where scrambling is abolished or reduced. Our work provides insights into the molecular mechanisms of hXkr4 scrambling and suggests the ability to thin membranes might be a general property of active scramblases.

## Introduction

In resting eukaryotic cells, the composition of the plasma membrane (PM) leaflets is [1–3]. The outer leaflet is primarily composed of phosphatidylcholine (PC) and sphingomyelin (SM) whereas the inner leaflet contains the negatively charged lipid phosphatidylserine (PS), phosphatidylethanolamine (PE), and phosphatidylinositols (PI’s, PIP’s) [1–3]. This asymmetry is generated by the activity of flippases and floppases, ATP-driven and lipid-specific pumps that respectively belong to the P-type ATPase and ABC transporter superfamilies and is essential for cellular homeostasis and membrane integrity. Phospholipid scramblases catalyze the rapid, non-specific, and bi-directional translocation of phospholipids between the two leaflets. At the PM, activation of scramblases causes loss of compositional asymmetry and results in the externalization of the signaling PE and PS lipids on the cell surface, which is a key trigger in multiple physiological processes, such as blood coagulation, membrane fusion or repair, and apoptosis [1–4].

Apoptosis is a highly organized and tightly regulated process where the activation of caspases leads to morphological changes of cells such as shrinkage, DNA fragmentation, blebbing, PS externalization, and cell death[5–7]. Apoptotic cells and their released fragments are identified and cleared by macrophages via dedicated PS receptors in a process called efferocytosis [8, 9]. Failure of efferocytosis, which can be caused by impaired PS externalization, leads to necrosis, where the release of intracellular components incites inflammatory and immunogenic reactions, which can lead to autoimmune responses or other pathological states [10–12].

The family of X Kell-related (Xkr) membrane proteins are evolutionarily conserved from nematodes to humans, and the human genome encodes for 9 homologues, Xkr1-9 [13]. Three human homologues, Xkr4, Xkr8, and Xkr9, and CED-8 from the nematode *Caenorhabditis elegans* were shown to mediate apoptotic scrambling in cells [14, 15]. Mutations and/or deletion of Xkr genes contribute to autoimmune disorders, such as systemic lupus erythematosus, favor inflammation, asthma, and lung cancer [16–21], further highlighting their importance in human physiology. Consistent with their broad physiological importance, the localization of Xkrs is variable: whereas Xkr8 is ubiquitously expressed, Xkr9 is predominantly expressed in the intestine, and Xkr4 localizes to the brain, nervous system, and eyes [14, 15, 22]. Mutations in Xkr4 affect cerebellar development [23], and have been implicated in neurological disorders such as Attention-Deficit/Hyperactivity Disorder (ADHD) [24] and substance abuse [25].

During apoptosis, the effector caspases, Casp3 in mammalian cells and CED-3 in *C. elegans* [7], cleave Xkr scramblases at a C- (in Xkr4, -8 and -9) [14, 15] or N-terminal site (in CED-8) [14, 26] to activate them and enable scrambling (Fig. 1 Supp. 1a). In Xkr8, constitutive, caspase independent scrambling is enabled by phosphorylation at three C-terminal residues [27], suggesting that cleavage is not strictly required for activation. It has been proposed that, following activation, Xkr4 and -8 oligomerize to scramble lipids [22, 28] (Fig. 1 Supp. 1b). Additionally, it has been suggested that Xkr4 activation also requires binding of a peptide from the nuclear DNA repair protein XRCC4 [22] and of extracellular Ca^2+^ [29]. However, full length or processed Xkr8 and -9 purify as monomers, and no scrambling activity was detected on their reconstitution in proteoliposomes [30, 31]. Further, the Xkr1 homologue, which lacks a caspase recognition site [15], functions in complex with VPS13 [32–34], and scrambles lipids when purified and reconstituted in liposomes [35].

**Figure 1.**
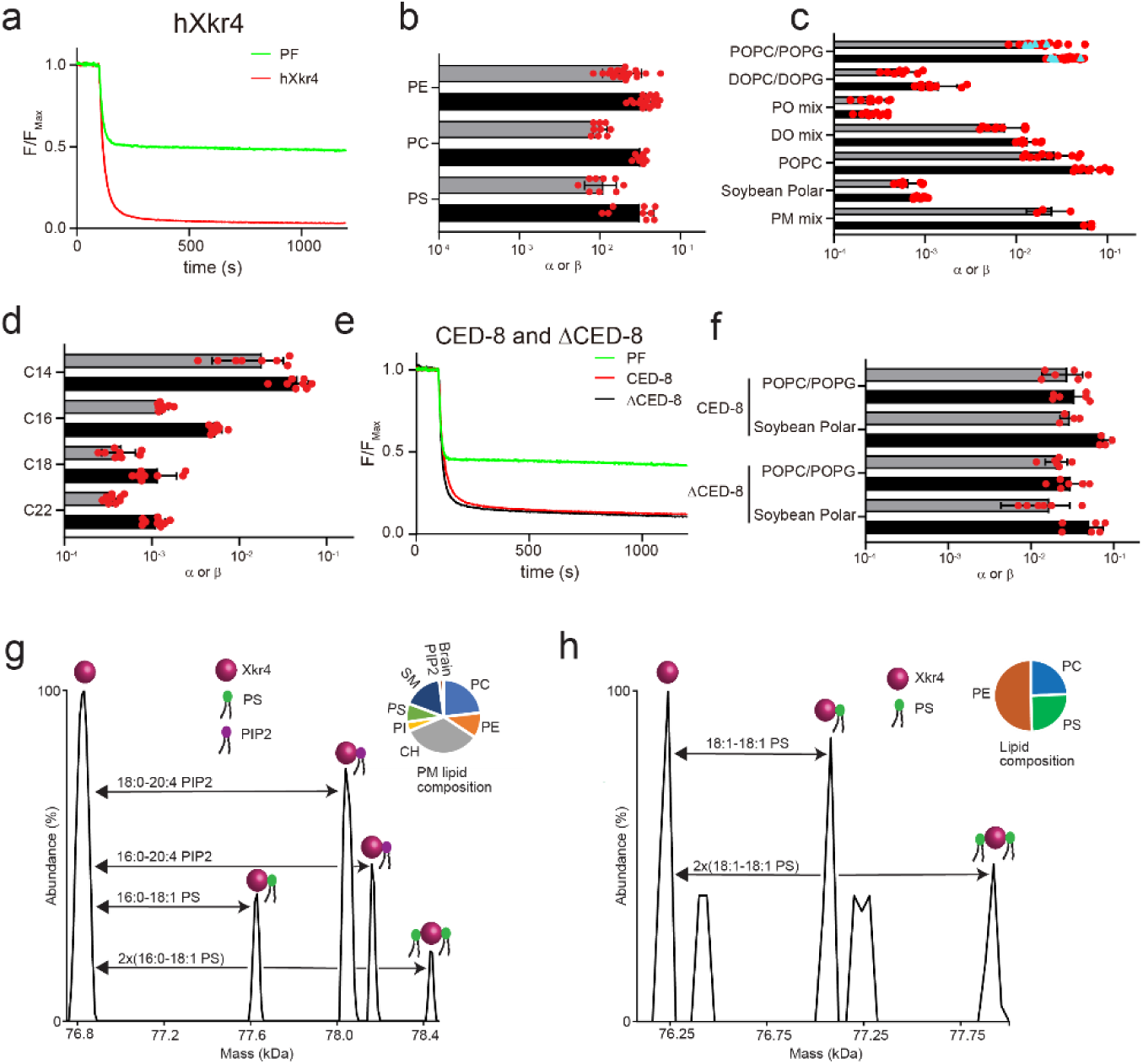
Characterization of hXkr4 and CED-8 in proteoliposomes. **a)** Representative traces of the dithionite induced fluorescence decay in the scrambling assay for protein free liposomes (green) and hXkr4 (red) reconstituted in 7:3 POPC:POPG mixed membranes. b) Forward (α) and reverse (β) scrambling rate constants of hXk4 reconstituted in 7:3 POPC:POPG mixed membranes doped with NBD-labeled PE, PC, or PS lipids. c-d) Forward (α) and reverse (β) scrambling rate constants of hXk4 reconstituted in membranes with different composition (c) or with fixed 7 PC: 3 PG headgroup and different acyl chain length (d). e) Representative traces of the dithionite induced fluorescence decay in the scrambling assay for protein free liposomes (green), CED-8 (red), and ΔCED-8 (black) reconstituted in 7:3 POPC:POPG mixed membranes. f) Forward (α) and reverse (β) scrambling rate constants of CED-8 and ΔCED-8 in liposomes formed from 7:3 POPC:POPG or Soybean Polar lipids. Bars in panels (b, c, d, f) are averages for α (black) and β (gray) (N ≥ 3), error bars are S. Dev., and red circles are values from individual repeats. g-h) g) Deconvoluted mass plot obtained from nMS analysis of hXkr4 from PM-mimicking liposomes (g) or from 2:1:1 DOPE:DOPC:DOPS (DO mix) liposomes. The relative lipid composition is given in the pie charts (insets) and in Supplementary Table 1.

The architecture of Xkr proteins was revealed by the recent cryoEM structures of detergent solubilized Xkr9 from *Rattus norvegicus* (rXkr9, PDBID: 7P14) [31] and of human Xkr8 (hXkr8, PDBID: 8XEJ) in complex with its ancillary subunit Basigin in detergent micelles and nanodiscs [30, 36]. Both structures were determined with the aid of antibodies to facilitate cryoEM imaging. We will use the hXkr8 as our reference since this homologue has been functionally characterized in greater detail, and the structures are very similar (Cα r.m.s.d. ∼1.38 Å). The Xkrs are comprised of 8 transmembrane helices (TM1-8) and 3 reentrant helices (IH1-3) arranged in two internal repeats of 4 TM helices and 1 hairpin (termed ND and CD, Fig. 1 Supp. 1c). Two hydrophobic, lipid filled, cavities, termed C1 and C2, are formed at the interface between the ND and CD repeats (Fig. 1 Supp. 1d, e) [30, 31]. The C1 cavity is constricted at the intracellular side by TM2, IH3 and by the C-terminal helix (Fig. 1 Supp. 1d). The C2 cavity, located on the opposite side of the protein, is hydrophobic and shallow (Fig. 1 Supp. 1e). The TM1 and TM3 in the ND contain several conserved polar and charged residues (Fig. 1 Supp. 1f). It has been proposed that upon activation of hXkr8, the TM1 and TM3 separate exposing these hydrophilic residues to the membrane core so that they could form a stairway for the lipid headgroups to move between leaflets [30], in a mechanism reminiscent of the credit card model of scrambling [37]. However, in the hXkr8 and rXkr9 cryoEM structures, these residues are isolated from the membrane by the close juxtaposition of TM1 and TM2s (Fig. 1 Supp. 1f) and cannot directly interact with lipids. No conformational rearrangements were seen in caspase processed rXkr9, besides the lack of the cleaved C-terminal helix [31], suggesting the known Xkr conformation might represent an inactive state. These residues were also shown to play a role in scrambling by hXkr4 [29]. However, their proposed role was to form a Ca^2+^ binding site whose occupancy prevents dynamic rearrangements of the TM1 and TM3 helices of the ND repeat, a process inferred to facilitate scrambling [29].

To gain insights into the basis of apoptotic scrambling by an active Xkr protein, we purified and functionally reconstituted the human Xkr4 (hXkr4) and CED-8 from *C. elegans*, which mediate apoptotic scrambling in cells [14, 15]. Both purified proteins mediate lipid scrambling when reconstituted in proteoliposomes, with properties that are modulated by physico-chemical properties of the membranes, such as thickness and rigidity, as expected for scramblases [38, 39]. Unexpectedly, we found that full-length hXkr4 and CED-8 scramble lipids and a construct corresponding to the N-terminally processed CED-8 is also active with properties similar to those of the wildtype protein. Using lipid vesicle native mass spectrometry (nMS) [40, 41] we show that full length hXkr4 is a monomer in liposomes and binds to the acidic phospholipids PS and PIP2. Thus, neither caspase cleavage nor oligomerization is required for function. We used cryoEM to determine the structure of hXkr4 alone and found that this active scramblase adopts a novel conformation, where the occlusion of the C1 cavity is relieved by a rotation of the ND and CD repeats. The ND also undergoes internal rearrangements which result in opening of a vestibule to the extracellular solution and in an altered electrostatic profile at the protein-membrane interface. Molecular dynamics (MD) simulations show that hXkr4 in the cryoEM conformation distorts and thins the membrane at the ND vestibule. This membrane thinning is more pronounced in lipid compositions where scrambling activity is favored, and is not seen in simulations of an AlphaFold2 [42] model of hXkr4 in an hXkr8-like conformation with a closed C1 cavity and ND vestibule. In silico mutagenesis experiments support the notion that the charged stairway residues in the ND vestibule play a role in scrambling. Our results reveal a novel conformation of the Xkr4 apoptotic scramblase and provide insight into their scrambling mechanism.

## Results

### Full length hXkr4 scrambles lipids

To test whether Xkr proteins are scramblases we sought to purify and functionally reconstitute them in proteoliposomes. A screen of GFP-tagged family members using fluorescence size exclusion chromatography (FSEC) [43, 44] identified human Xkr4 (hXkr4, MW ∼71kDa) as a promising candidate (Fig. 1 Supp. 2a). hXkr4 mediates apoptotic scrambling in cells [15, 22, 29] and has a C-terminal caspase cleavage site distal from the membrane (Fig. 1 Supp. 1a). On a calibrated size exclusion chromatography column, purified full length hXkr4 in 0.05% (w/v) dodecyl-β-D-maltoside (DDM)- 0.01% (w/v) cholesteryl hemisuccinate (CHS) and 0.001% lauryl maltose neopentyl glycol (LMNG)- 0.0001% CHS elutes with a main peak at an elution volume consistent with a monomer (Fig. 2 Supp. 1a). We used a well-characterized in vitro assay [45, 46] to determine whether hXkr4 is a lipid scramblase (Fig. 1 Supp. 2b). Briefly, proteoliposomes reconstituted with trace amounts of acyl-chain labeled NBD-phospholipids (NBD-PLs) are treated with the membrane-impermeant, reducing agent dithionite which can access and irreversibly reduce only NBD fluorophores in the extraliposomal leaflet (Fig. 1 Supp. 2b). Therefore, in protein-free liposomes (Fig. 1a, Fig. 1 Supp. 2c) or in proteoliposomes with a non-scramblase protein, such as the CLC-ec1 exchanger (Fig. 1 Supp. 2d), only ∼50% reduction in fluorescence is seen. In proteoliposomes containing an active scramblase a more pronounced fluorescence loss is observed as inner leaflet labeled lipids are scrambled to the outer leaflet (Fig. 1 Supp. 2b) [45, 46]. We reconstituted hXkr4 in proteoliposomes formed from a 7:3 mixture of 1-Palmitoyl-2-oleoyl-sn-glycero-3-phosphocholine/ 1-Palmitoyl-2-oleoyl-sn-glycero-3-[6 phospho-rac-(1-glycerol)] (POPC/POPG). Addition of dithionite leads to a pronounced fluorescence loss which reaches ∼75- 80% at steady state (Fig. 1a) with macroscopic scrambling rate constants of ∼3.9×10^-2^ s^-1^ (Fig. 1b-c), which are comparable to those of the nhTMEM16 and afTMEM16 scramblases in the presence of Ca^2+^ [45, 47]. Similar results were obtained using a BSA back-extraction assay [48] (Fig. 1 Supp. 2e, f), indicating that reconstituted hXkr4 does not allow entry of dithionite into the liposomes by mediating ion transport or by destabilizing the membrane. Thus, purified full-length hXkr4 is a lipid scramblase.

**Figure 2.**
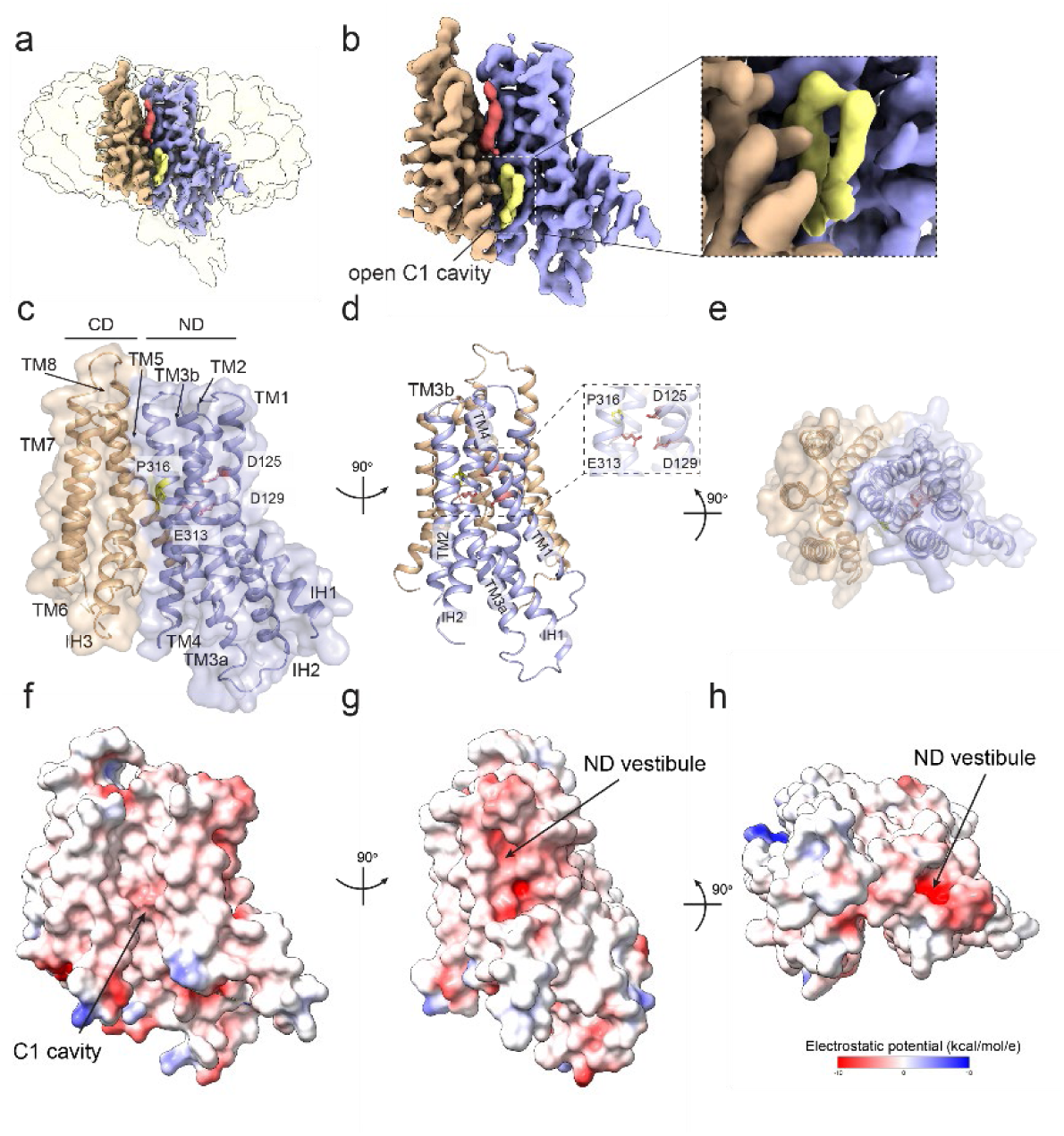
Structure of hXkr4. **(a-b)** CryoEM density of hXkr4 in LMNG-CHS detergent micelles. The ND repeat is colored in pale blue, the CD repeat in wheat. Associated lipid-like densities are shown in yellow (inner leaflet) and red (outer leaflet). Inset shows a close-up view of the lipid-like density in the C1 cavity. c-e) The structure of hXkr4 viewed from the plane of the membrane (c), from the side of the ND repeat (d), and from the extracellular solution (e). The protein is shown in ribbon representation with the ND repeat in pale blue, the CD repeat in wheat, the charged stairway residues (D125, D129, and E313 in pink) and P316 (in yellow) are shown in stick representation. The transparent surface representation of the protein is shown in (c) and (e). f-h) Electrostatic potential plotted on the surface of hXkr4 from the same views as in (c-e).

### Caspase cleavage is not required for the *in vitro* activity of Xkr4 and CED-8

To investigate the role of caspase cleavage in scrambling mediated by purified Xkr proteins we incubated hXkr4 with purified CASP3 [49]. However, the processed protein was unstable and could not be functionally reconstituted. Similarly, expression of a construct corresponding to the caspase processed hXkr4 (residues 1-564) was insufficient for functional analyses. To circumvent these limitations, we purified full-length CED-8, the *C. elegans* Xkr homologue, and ΔCED-8 which lacks the first 21 N-terminal residues and corresponds to the caspase processed construct [14, 26]. Reconstitution of full-length CED-8 and of ΔCED-8 in liposomes formed from soybean polar, or POPC/POPG lipids shows both constructs are active scramblases with similar scrambling rate constants (∼4×10^-2^ s^-1^) in both lipid compositions (Fig. 1e, f), which are comparable to those of hXkr4 (Fig. 1c). These results show that caspase cleavage is not required for the *in vitro* scrambling activity of hXkr4 and CED-8. While surprising, these findings are consistent with reports showing that purified full-length Xkr1 also scrambles lipids [35] and that Xkr8 can mediate caspase-independent scrambling in cells [27].

### Modulation of hXkr4 scrambling activity by membrane properties

The activity of many scramblases, such as the TMEM16s and GPCRs, is characterized by poor selectivity for the headgroups of the transported lipids and by a sensitivity to changes in membrane properties [38, 39, 45, 47, 48, 50–52]. Therefore, we tested whether hXkr4 shares these characteristics. We found that it scrambles tail-labeled PE, PC and PS lipids with similar rate constants (Fig. 1b, Fig. 1 Supp. 2c), indicative of poor headgroup selectivity. Then we measured its scrambling rate constants in liposomes formed from the following membrane compositions: 100% POPC, two simple headgroup mixtures, 7:3 PC:PG and 2:1:1 PE:PC:PS lipids with DO or PO acyl tails (referred to as DO-mix and PO-mix), a complex mixture mimicking the composition of the plasma membrane (referred to as PM-like) [40, 41] (Supplementary Table 1), and soybean polar lipid extract (Fig. 1c, Fig. 1 Supp. 2c). hXkr4 activity is maximal in pure POPC, 7:3 POPC:POPG or PM-like liposomes (α, β∼3.2-7.0×10^-2^ s^-1^), is intermediate in vesicles formed from DO-mix and 7 DOPC: 3 DOPG (α, β∼1-10×10^-3^ s^-1^), and is nearly ablated in Soybean polar and PO-mix lipids (α, β<10^-3^ s^-1^) (Fig. 1c). Next, we tested how changes in membrane thickness from ∼32 to ∼41 Å [38] affect hXkr4 scrambling by systematically changing the acyl chain length, from 14 to 22 carbons (C14 to C22), of the 7:3 PC:PG mix. We found that scrambling is maximal in C14 lipids, slightly slower in C16 lipids, and is reduced ∼30-fold in the C18 and C22 lipids (Fig. 1d, Fig. 1 Supp. 2c). Thus, like other scramblases, hXkr4 does not select among transported lipid headgroups, is impaired in thicker membranes, and by POPE-containing membranes that facilitate the formation of liquid-ordered domains. These results show that membrane composition is a critical regulator of hXkr4 function, but that no specific effects can be ascribed to lipid acyl chain saturation or headgroup composition. These functional properties closely mirror those of TMEM16 scramblases [38, 47, 48].

### Monomeric hXkr4 is a functional scramblase

Our experiments show that hXkr4 purifies as a monomer (Fig. 1 Supp. 2a, Fig. 2 Supp. 1a) and is an active scramblase (Fig. 1a-d). To test whether active hXkr4 adopts a different oligomeric state in membranes, as recently proposed [22, 28], we employed the recently developed lipid vesicle native mass spectrometry (nMS) approach [40, 41] to determine the mass of hXkr4 reconstituted in PM-like or DO-mix liposomes, where the protein is active (Fig. 1c) (Supplementary Table 1). In both lipid compositions the major peak in the spectra corresponds to the mass of full-length monomeric Xkr4 (Fig. 1g, h). Interestingly, in both cases we detect multiple peaks with MW shifts matching those of 1 or 2 bound PS lipids (Fig. 1g, h). Further, in PM-like liposomes we also observe two additional peaks indicating that the two major brain PIP2 species, 18:0-20:4 and 16:0-20:4 PIP2, can also bind to purified hXkr4 (Fig. 1g). In neither case a peak corresponding to higher order oligomers was visible. Therefore, monomeric hXkr4 is a functional scramblase.

### CryoEM structure of hXkr4

We used cryoEM imaging to understand the structural basis of scrambling by the purified and active monomeric hXkr4 scramblase. We chose to image the protein alone, to avoid potential conformational biasing from binding of antibodies. Initial imaging experiments of hXkr4 solubilized in DDM-CHS were unsuccessful, likely reflecting the small size of the protein (∼71 kDa) and the presence of excess empty micelles that lower the protein signal. Inspired by recent work on the GAT1 transporter [53], we imaged hXkr4 solubilized in the low CMC detergent LMNG-CHS at sub-CMC nominal concentration to minimize the number of empty micelles. We collected ∼25,000 micrographs from regions with thin ice (majority with 15-40 nm thickness) (Fig. 2 Supp. 1). Extensive data processing in cryoSPARC [54] resulted in a sharpened map with an average resolution of 3.72 Å (Fig. 2a, b, Fig. 2 Supp. 1c-f, Table 1) and enabled building of the atomic model for the transmembrane region of hXkr4 (Fig. 2c-e, Fig. 2 Supp. 1i). Density for the cytosolic N- and C-termini is poor, suggesting these regions are flexible and dynamic in the cytosol (Fig. 2a, Fig. 2 Supp. 1h). The overall fold of hXkr4 resembles that of hXkr8 and rXkr9, with 8 TM helices (TM1-8) and 3 short intramembrane helices (IH1-3) (Fig. 2c). The ND and CD repeats are related by pseudo-2-fold symmetry, and respectively consist of TMs 1-4, IH1, and IH2, and TMs 5-8 and IH3 (Fig. 2c). The TM3 helix is broken around P316 into two short helices, TM3a and TM3b, connected by a short intramembrane loop which contains the negatively charged side chain of E313 (Fig. 2c-e).

**Table 1.** Cryo-EM data collection, refinement, and validation statistics.

|  |  |
| --- | --- |
|  | Human Xkr4<br>(EMDB-44744)<br>(PDB 9BOJ) |
| **Data collection and processing** |  |
| Magnification | 105000 |
| Voltage (kV) | 300 |
| Electron exposure (e-/Å<sup>2</sup>) | 58.28 |
| Defocus range (µm) | 0.9-2.6 |
| Pixel size (Å) | 0.825 |
| Symmetry imposed | C1 |
| Initial particle images (no.) | 5,540,840 |
| Final particle images (no.) | 446,787 |
| Map resolution (Å) | 3.72 |
| FSC threshold | 0.143 |
| Map resolution range (Å) | 3.0-4.6 |
| **Refinement** |  |
| Initial model used (PDB code) | N/A |
| Model resolution (Å) | 3.7 |
| FSC threshold | 0.143 |
| Model resolution range (Å) | 3.1-3.9 |
| Map sharpening *B* factor (Å<sup>2</sup>) | -150 |
| Model composition |  |
| Non-hydrogen atoms | 2551 |
| Protein residues | 321 |
| Ligands | 0 |
| *B* factors (Å<sup>2</sup>) |  |
| Protein | 55.42/118.27/78.48(min/max/mean) |
| Ligand | N/A |
| R.m.s. deviations |  |
| Bond lengths (Å) | 0.002 |
| Bond angles (°) | 0.442 |
| Validation |  |
| MolProbity score | 2.06 |
| Clashscore | 4.20 |
| Poor rotamers (%) | 3.57 |
| Ramachandran plot |  |
| Favored (%) | 93.33 |
| Allowed (%) | 6.67 |
| Disallowed (%) | 0 |

In our structure, TM2 is separated from IH3 so that the C1 cavity is wide-open to the hydrocarbon core of the bilayer (Fig. 2a-c, Fig. 2 Supp. 2a), suggesting it can accommodate lipids. Indeed, a non-protein density with lipid-like features with two connected tails is visible in its intracellular vestibule (Fig. 2b, inset). We also observe a weak elongated density in the extracellular region of this cavity (Fig. 2b), in the same region where a lipid was observed in the Xkr8 and 9 structures (Fig. 1 Supp. 1d) [30, 31]. The interior of the opened C1 cavity is hydrophobic and lined by the TM2, 3, 4, and 6 helices (Fig. 2c, f). The C2 cavity is shallow, exposed to the membrane, and hydrophobic (Fig. 2 Supp. 2b). Interestingly, there is a deep vestibule within the ND repeat that is directly exposed to the extracellular solution with the negatively charged residues D125, D129, and E313 at its deepest point (Fig. 2d-e, h). These residues correspond to the stairway residues identified in hXkr8 [30] (Fig. 1 Supp. 1f). Although these residues are not directly exposed to the bilayer, the membrane-exposed portion of ND vestibule is strongly electronegative (Fig. 2g). Since purified hXkr4 is a functional and monomeric scramblase, we hypothesize this conformation corresponds to a scrambling-competent state of the protein.

The present hXkr4 conformation presents notable differences from that adopted by hXkr8 and rXkr9 [30, 31]. The major rearrangement is a rotation of the ND and CD internal repeats which results in the opening of the C1 cavity (Fig. 3a, b). An alignment of hXkr4 to hXkr8 on their respective CD’s shows that the ND of hXkr4 is rotated relative to that of hXkr8 (Fig. 3a). This movement displaces the TM2 helix in the ND from the IH3 in the CD relieving the constriction that occludes the C1 cavity at its intracellular vestibule in hXkr8 (Fig. 3a). In hXkr8, this vestibule is occupied by the short C-terminal helix which interacts with the cytosolic portions of TM2, TM3 and TM4 from the ND repeat and of TM5, IH3, and TM7 from the CD (Fig. 3 Supp. 1a). In contrast, the weak density for the C-terminus in our hXkr4 map indicates this region is dynamic and in the cytosolic milieu (Fig. 2 Supp1h). However, the hydrophobic character of the opened C1 cavity interior renders it poorly suited to serve as a scrambling pathway for the hydrophilic lipid headgroups (Fig. 2f). Indeed, the density for lipids observed in this region suggest they are perpendicular to the membrane plane, suggesting the bilayer is unperturbed in this region (Fig. 1 Supp. 1d, Fig. 2b, inset). This is unlike the pronounced membrane thinning and severely tilted lipid orientations that enable scrambling by the TMEM16s [38, 47]. To investigate whether the C1 cavity is the scrambling pathway we introduced two bulky tryptophan side chains at two heights within the membrane: one demarcated at L147 on TM2 and G402 on TM6, and at S158 on TM2 and V436 on IH3. If the C1 cavity serves as the lipid pathway, then we expect that the constrictions caused by the bulky Trp side chains should impair lipid scrambling. We found that both double mutants, L147W G402W and S158W V436W, have WT-like activity when reconstituted in DO- mix vesicles (Fig. 3d), suggesting that the C1 cavity is unlikely to serve as the lipid pathway.

**Figure 3.**
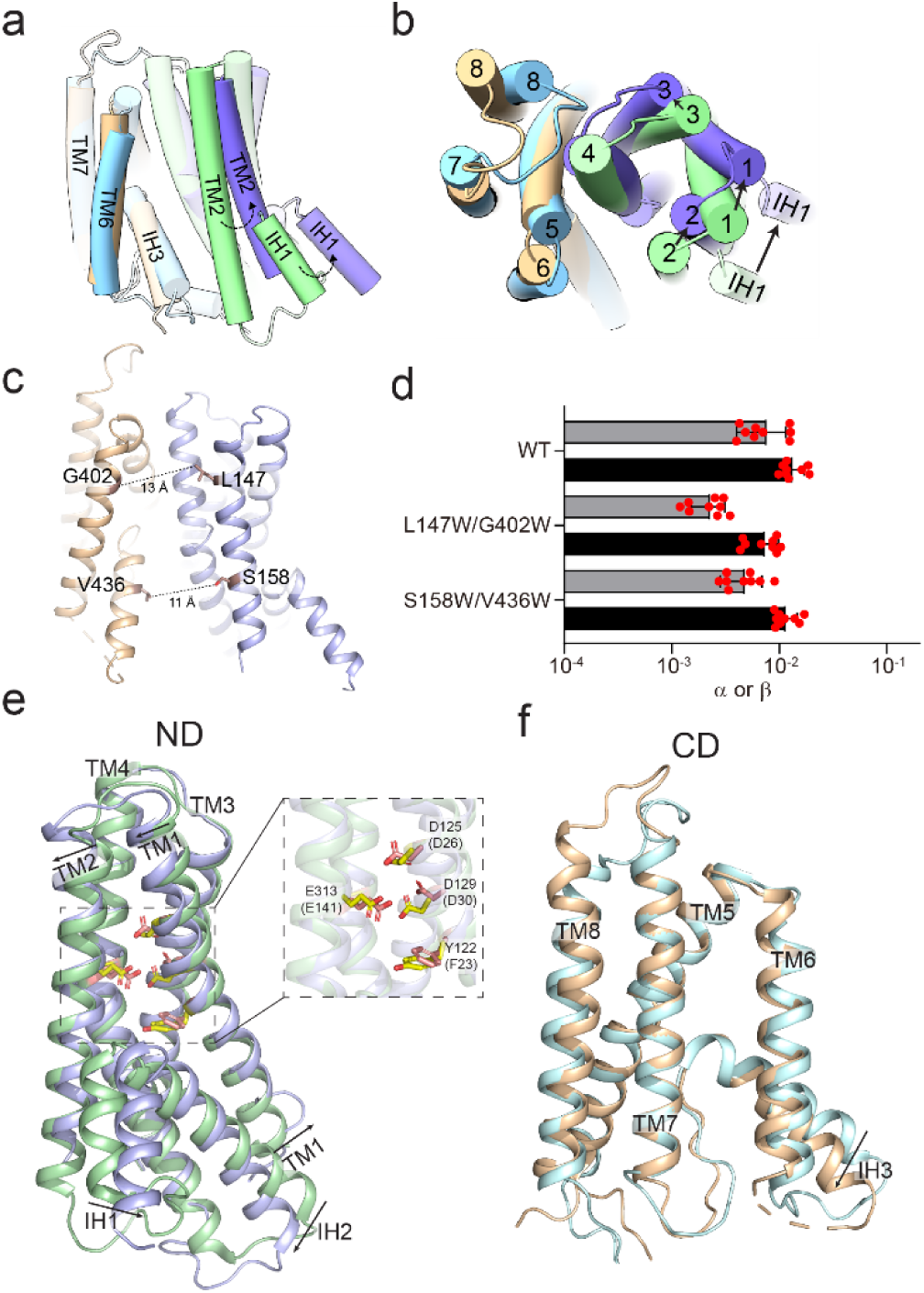
Structural changes in Xkr4. **a-b)** The cryoEM structures of hXkr4 and hXkr8 (PDBID: 8XEJ), shown in cylindrical cartoon representations, are aligned on their respective CD domains (wheat for hXkr4 and cyan for hXkr8). The pseudo-symmetry axis (dashed vertical line, panel a) and angle of rotation of the ND of hXkr4 (pale blue) relative to the ND of hXkr8 (pale green) is viewed from the plane of the membrane (a) or from the extracellular solution (b). The C-terminal helix of hXkr8 is colored in pink. c) The distance between the Cα atoms (maroon spheres) of L147 and S158 on TM2 and of G402W on TM6 and V436 on IH3 is shown (dashed lines). d) Forward (α) and reverse (β) scrambling rate constants of WT, L147W/G402W, S158W/V436W hXkr4 reconstituted in DO-Mix liposomes. Bars are averages for α (black) and β (gray) (N ≥ 3), error bars are S. Dev., and red circles are values from individual repeats. e-f) Alignment of the ND (e) and CD (f) repeats of hXkr4. Colors as in (a-b). Arrows denote direction of movement of the helices from hXkr8 to hXkr4. The charged residues in the ND of hXkr4 (hXkr8), D125 (D26), D129 (D30), E313 (E141), and Y122 (F23) are shown in stick representation and colored in yellow CPK (hXkr4) or pink CPK (hXkr8). Inset of (e) shows a close-up view of these residues.

The second major difference in the hXkr4 conformation is that the ND (Fig. 3e), but not the CD, undergoes significant internal rearrangements (Fig. 3f). An alignment of the individual internal repeats of hXkr4 and -8 shows the Cα r.m.s.d. of the ND’s is ∼1.9 Å and of the CD’s is ∼0.9 Å (Fig. 3e, f). The difference in the ND’s is due to a tilting of the TM1 helix around Y122, a reorientation of TM2, and a lateral displacement of the IH1 and IH2 helices (Fig. 3e). These rearrangements are enabled by the reduction in the interaction surface between the ND and CD repeats. To quantify this change, we used AlphaFold2 [42] to generate a model of hXkr4, hXkr4^α^, in an Xkr-8 like conformation with a closed C1 cavity (Cα r.m.s.d. ∼ 1.8 Å to Xkr8) (Fig. 3 Supp. 1c). When the C1 cavity is closed, the inter-repeat surface is primarily mediated by TM3 and TM4, with minor contributions from TM1 and TM2 (Fig. 3 Supp. 1d-e). When the C1 cavity opens, the TM2 and TM3a helices lose their interactions with the CD repeat (Fig. 3 Supp. 1d, f; Supp. Movie 1), allowing their rearrangements. In Xkr8 the ND repeat vestibule is closed to the extracellular solution and to the membrane, so that the electrostatic profile of the ND is nearly neutral (Fig. 3 Supp. 1b). Thus, the slight rearrangements in the TM1 and TM2 helices affect the electrostatic profile of the ND vestibule which is determined by the charged stairway residues (Fig. 2f-h, Fig. 3 Supp. 1b; Supp. Movie 2). Indeed, whereas the position of D125 (D26 in Xkr8) and E313 (E141 in Xkr8) is similar in the two structures, the rearrangement in TM1 displaces the side chain of D129 (D30 in Xkr8) so that it is closer to the membrane interface (Fig. 3e inset). These residues are conserved between hXkr4 (Fig. 2d) and Xkr8 (Fig. 1 Supp. 1e) and are important for lipid scrambling by the latter [30]. Therefore, we hypothesize that the rearrangements in the ND repeat which alter their exposure to the membrane might underlie the activity of hXkr4.

### Hydration and membrane thinning by the active hXkr4 conformation

We used molecular dynamics (MD) simulations to investigate how hXkr4 interacts with the membrane lipids. To mimic our experimental conditions, we simulated hXkr4 in 100 mM KCl and using two membrane compositions: 100% POPC lipids, where the protein is maximally active, and DO-mixed membranes, in which the protein is moderately active (Fig. 1C). For each system considered we quantitatively analyzed 10 independent replicas of 500 ns long trajectories and examined dynamic rearrangements in the protein, the protein-lipid interface, ion binding, and hydration state (Fig. 4, Supplementary Table 2). For WT hXkr4 in POPC membranes we ran 10 replicas using GROMACS and 10 using AMBER, with no significant differences (Fig. 4 Supp. 1a-e). Therefore, we considered 20 total replicas for this condition.

**Figure 4.**
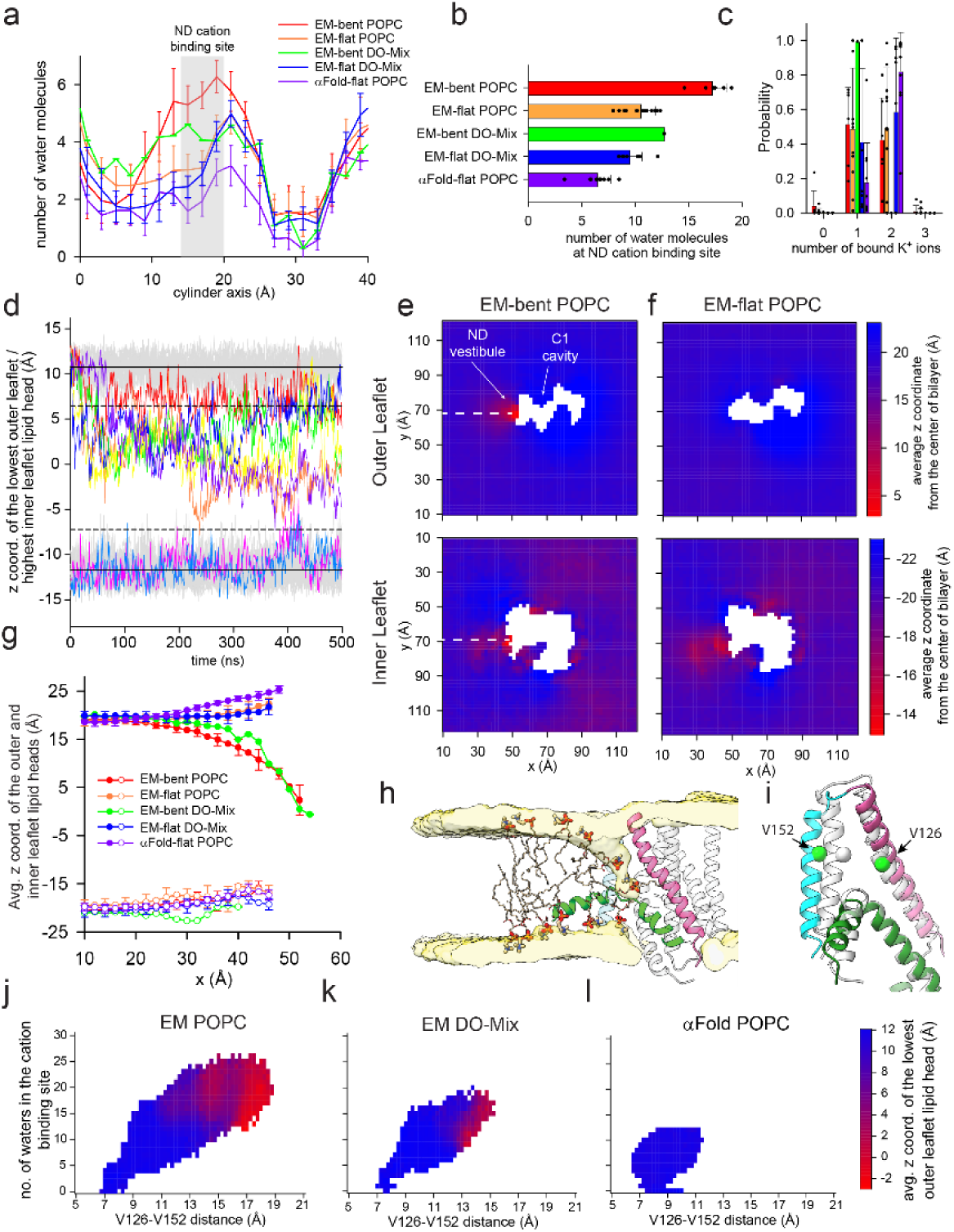
Hydration, ion binding and membrane thinning by hXkr4. **a)** The average number of water molecules along the cylindrical axis along the ND vestibule (Fig. 4 Supp. 1a) for trajectories of cryoEM hXkr4 where the membrane bends (EM-bent) or where it remains flat (EM-flat), in POPC lipids (EM-bent POPC, red, n=6; EM-flat POPC, orange, n=14), or in DO-Mix membranes (EM-bent DO-Mix, green, n=1; EM-flat DO-Mix, blue, n=9), and of αFold model hXkr4 in POPC (αFold-flat POPC, purple, n=10). See Methods for details. The region of the cation site in the ND vestibule (near D125, D129, and E313, 16 Å<h<20 Å) is colored in gray. (b-c) The average total number of water molecules in the ND cation site (b) and the probability distribution of the number of K^+^ ions in the ND cation site (c) for the same trajectory groups as in (a). Filled black circles in (b-c) represent values from individual trajectories. Error bars in (a-c) are the St.Dev. of the values from individual trajectories. For EM-bent DO-Mix n=1, as we observe membrane bending only in 1 trajectory, so no error is reported. (d) Time evolution of the z coordinate of the phosphorous atom in the lowest/highest outer/inner leaflet lipid headgroup of individual trajectories of EM-flat POPC (gray) and EM-bent POPC (colored). Solid and dotted lines represent the Avg. and Avg.+ or - 3xSt.Dev of the z coordinate for IL and OL, respectively. (e-f) Two-dimensional (2D) plot of the average z coordinate of the phosphorous atoms in the outer (top panel) and inner (bottom panel) leaflet lipid headgroups on the x-y plane of the simulation box, calculated from EM-bent POPC (e) or EM-flat POPC (f) trajectories. Individual pixels are colored from red to blue by the average z coordinate displacement. (g) Cross-section of the 2D plot calculated along the white dotted lines in (e-f) for the outer (filled circles) and inner (empty circles) leaflets. Data is Mean± St.Dev. (h) Representative snapshot from a EM-bent POPC trajectory. hXkr4 is shown in cartoon representation with TM1 (pink), TM2 (cyan), and IH1 (green), TM3-8 are in light gray. The average lipid head density of the outer and inner leaflets is shown in surface representation (light yellow). Representative lipid molecules are shown in stick, headgroup atoms in thicker sticks. (i) Superposition of the TM1, TM2, and IH1 from the cryoEM (light gray) and representative MD frame (colored as in h). Cα atoms of V126 in TM1 and V152 in TM2 are shown as sphere (green). (j-l) The average z coordinate of the phosphorous atom of the headgroups from the lowest outer leaflet lipid (colored from is plotted as a function of the V126-V152 Cα distance (x axis) and of the number of water molecules in the cation site in the ND vestibule (y axis) for trajectories for cryoEM hXkr4 in POPC (j) or DO-Mix membranes (j) or for αFold hXkr4 POPC (l).

Since membrane deformation and thinning are important for lipid flip-flop by other scramblases [38, 39, 47, 55–62], we inspected whether the membrane near the C1 cavity or around other regions of the cryoEM conformation of hXkr4 is perturbed in our simulations. In no trajectories we observe membrane deformation or water penetration near the hydrophobic C1 cavity (Fig. 4e-f, Fig. 4 Supp. 1a), consistent with the idea that this region is not the lipid scrambling pathway. In contrast, the open ND vestibule becomes hydrated within the first 20 ns and remains such throughout all trajectories (Fig. 4a, b, Fig. 4 Supp. 1f). We also observe that one or two K^+^ ions spontaneously enter the vestibule from the extracellular solution and interact with the negatively charged side chains of D125, D129, and E313 (Fig. 4c, Fig. 4 Supp. 1a). The residency time of individual K^+^ ions within the vestibule is low and ions frequently exchange between the extracellular milieu and the vestibule, suggesting that K^+^ binding is not stable. Notably, this region was proposed to serve as a Ca^2+^ binding site in Xkr4 [29], suggesting ion binding might be mainly driven by the negative electrostatic profile of this region (see below).

In 6 of 20 trajectories in POPC membranes we observe that outer leaflet (OL) lipids spontaneously rearrange near the ND vestibule (bent OL trajectories) (Fig. 4d-h), so that their headgroup approach the membrane-exposed charged stairway residues via the widened TM1-TM2 fenestration (Fig. 4h). In this region, lipids adopt tilted poses relative to the membrane plane, in some cases becoming nearly parallel to it (Fig. 4h). Finally, we also rarely observe a more modest deformation of the inner leaflet (IL) at the ND vestibule (in 2 of 20 trajectories), near the short intramembrane helical turn formed by IH1 and IH2 (Fig. 4d-h). In all cases the thinning deformation remains relatively local to the ND vestibule (Fig. 4e). Thus, near the ND vestibule there is a pronounced local thinning of the membrane, with the lipid headgroups from the outer and inner leaflets coming within ∼15 Å of each other (Fig. 4h). In DO mix bilayers, where hXkr4 has intermediate activity (Fig. 1c), we observe membrane thinning only in 1 of 10 trajectories (Fig. 4 Supp. 1g). Besides the difference in frequency of membrane bending, the trajectories in DO mix bilayers closely resemble the corresponding ones in POPC membranes in terms of vestibule hydration (Fig. 4a, b), average K^+^ occupancy (Fig. 4c), and extent of membrane bending in the OL and IL (Fig. 4g), indicating that membrane composition primarily affects the frequency of these membrane thinning events. A steric constriction defined by the IH1, IH2 and TM2 helices (Fig. 4h) prevents the lipid headgroups from freely moving between leaflets, suggesting additional rearrangements might be needed to allow scrambling. Notably, membrane bending correlates with the hydration state of the ND vestibule near the location of the charged stairway residues D125, D129, and E313: in the 6 bent OL trajectories this region is occupied by ∼16 water molecules, while in the remaining 14 flat OL trajectories the average water occupancy is ∼10 (Fig. 4a, b). In contrast, there is no difference in the K^+^ occupancy of the ND vestibule between the flat and bent OL trajectories (Fig. 4c). Thus, membrane thinning correlates with hydration of the ND vestibule. Our simulations show that rearrangements of the TM1 and TM2 helices underlie these different hydration states. Specifically, in POPC and DO mix trajectories we see that more pronounced membrane bending and hydration correlate with widening of the membrane fenestration into the ND vestibule, quantified by the distance between V126 on TM1 and V152 on TM2 (Fig. 4i-k, l), ∼8 and ∼9 Å in the starting conformations of hXkr4 and hXkr4^α^ simulations. When the two helices remain near the starting positions, the membrane is flat, and the vestibule is poorly hydrated (Fig. 4j-k). In contrast, as the helices become progressively more separated the membrane becomes more bent and the vestibule is more hydrated (Fig. 4j-k). These results suggest that dynamic openings of the ND vestibule promote its increased hydration and membrane thinning.

To test whether the increased dynamics of TM1 and TM2 are enabled by the cryoEM conformation of Xkr4 we simulated hXkr4^α^ in 100 mM KCl and POPC membranes, conditions in which membrane bending occurs more frequently for the cryoEM conformation. In the hXkr4^α^ trajectories the probability of double K^+^ occupancy of the ND vestibule is increased compared to that of the hXkr4 cryoEM conformation (Fig. 4c). In contrast, the water occupancy of the ND vestibule is reduced to ∼6 molecules (Fig. 4a, b), and we never observe bending of the OL (Fig. 4 Supp. 1h). Strikingly, the TM1 and TM2 helices in hXkr4^α^ are non-dynamic, as they sample only limited deviations from their original conformation (Fig. 4l). These observations are consistent with the lack of membrane thinning by Xkr8 reconstituted in nanodiscs [36]. Thus, the cryoEM conformation of the active hXkr4 scramblase allows the TM1 and TM2 helices to dynamically sample states with more open ND vestibule fenestration which in turn enable membrane thinning. The frequency and extent of membrane thinning in our simulations correlates with the degree of activity of the protein: thinning is most frequent in conditions of maximal scrambling activity (hXkr4 cryoEM conformation in POPC), is intermediate in conditions of medium activity (hXkr4 cryoEM conformation in DO-mix lipids), and not detected in conditions where the protein is inactive (hXkr4^α^, in a Xkr8-like conformation) (Fig. 4, Fig. 4 Supp. 1h). These observations suggest that that the membrane thinning facilitated by the exposure of the charged residues in ND vestibule is mechanistically related to lipid scrambling by hXkr4.

### Role of charged residues in membrane thinning and lipid scrambling

We tested this hypothesis by generating three single charge-neutralizing mutants, D125A, D129A, and E313A via in silico mutagenesis and simulating each construct in POPC membranes and 100 mM KCl in 10 replicas of 500 ns. In all mutants we observed lower K^+^ occupancy of the vestibule (Fig. 5a), consistent with the reduction in negative charge of the vestibule due to the mutation. The frequency of membrane bending is slightly reduced in the trajectories of the three mutants: it occurs in 2 of 10 trajectories of D125A, and in 1 of 10 for D129A and E313A (Fig. 5 Supp. 1a). The trajectories with flat or bent membranes were very similar to the corresponding ones of the WT protein in terms of vestibule hydration (Fig. 5b, c), membrane thinning (Fig. 5d), and sampling of the conformational space (Fig. 5e). These findings suggest that the overall electrostatic profile of the ND vestibule facilitates membrane thinning. We attempted to express and purify the three mutants of hXkr4, however their expression levels were too low for functional reconstitution. Importantly, these residues correspond to the stairway mutants that are critical for scrambling by hXkr8 in cells [30], supporting the idea these charged side chains play a key role in enabling Xkr scrambling.

**Figure 5.**
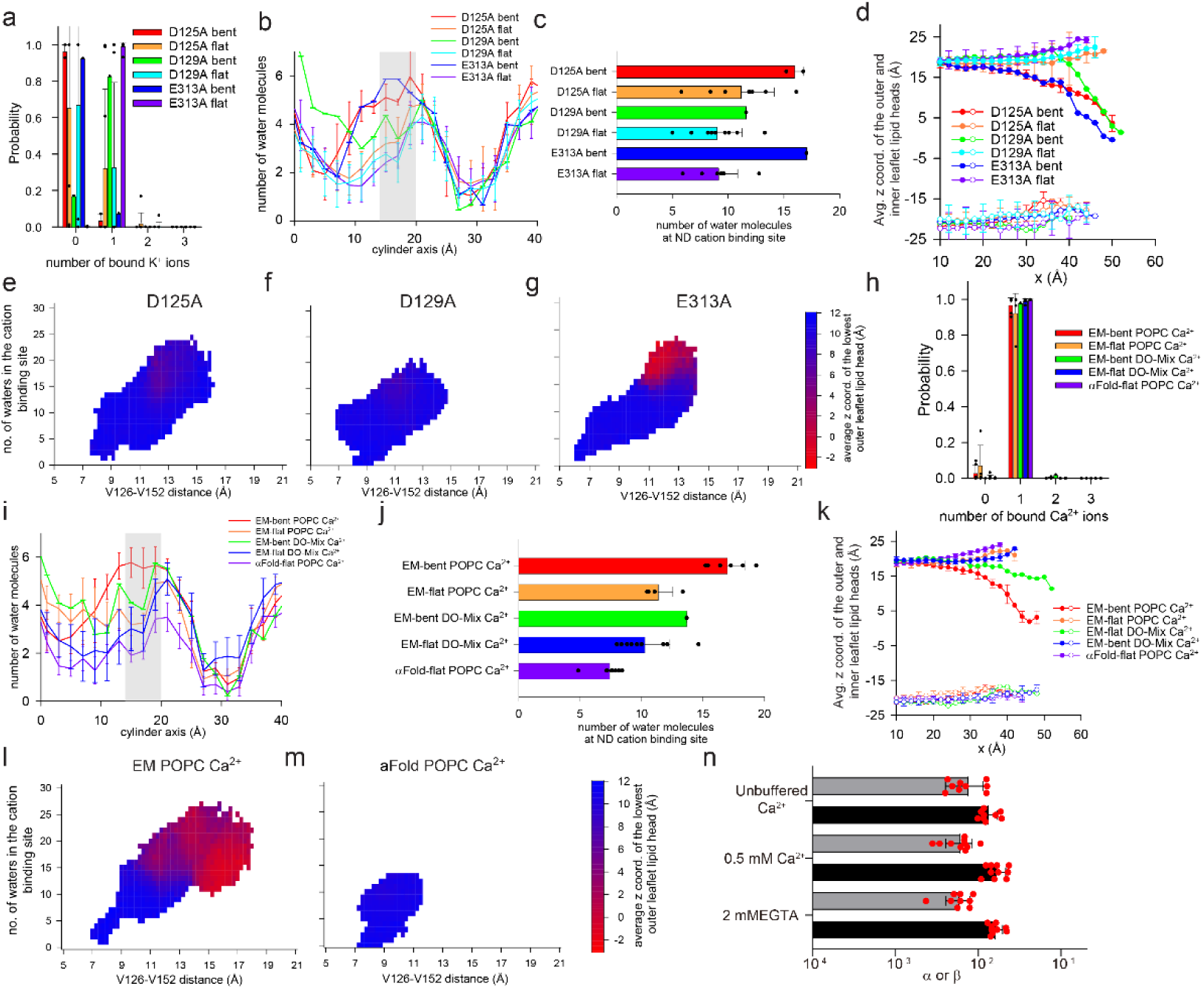
Role of charged stairway residues and Ca^2+^ binding to hXkr4. a-g) Probability distribution of the number of K^+^ ions in the ND cation site (a), distribution of water molecules in the ND repeat vestibule (b), total number of water molecules in the ND cation site (c), average z coordinate of the phosphorous atom of the lower/highest lipid headgroup from the outer/inner leaflet in MD trajectories of hXkr4 D125A bent (red), D125A flat (orange), D129A bent (green), D129A flat (cyan), E313 bent (blue), or E313 flat (purple) in POPC membranes (d). (e-g) the average z coordinate of the phosphorous atom of the headgroups from the lowest outer leaflet lipid (colored from is plotted as a function of the V126-V152 Cα distance (x axis) and of the number of water molecules in the cation site in the ND vestibule (y axis) for trajectories for D125A (e), D129A (f), and E313A (g). (h-m) same plots as in (a-g) but for simulations of hXkr4 in cryoEM and αFold conformations in POPC membranes and in the presence of Ca^2+^. n) Forward (α) and reverse (β) scrambling rate constants of hXk4 reconstituted in DO-Mix liposomes in unbuffered Ca^2+^ (∼10 µM), 2 mM EGTA (<10 nM Ca^2+^), and 0.5 mM Ca^2+^. Data in all panels is Mean± St.Dev.

### Cation binding to the opened vestibule

Recently, Ca^2+^ was proposed to bind to hXkr4 [29] in a location near where we observe spontaneous K^+^ binding in the ND vestibule (Fig. 4e, f). We investigated Ca^2+^ binding in 10 independent 500 ns long simulation trajectories of the cryoEM conformation of hXkr4 in POPC or DO mixed membranes and of hXkr4^α^ in POPC bilayers. We replaced the 100 mM KCl with 100 mM CaCl_2_, to compare identical concentrations of the two cations. In all trajectories a single Ca^2+^ enters the vestibule within the first 50 ns and remains stably bound throughout (Fig. 5h, Fig. 5 Supp. 1d). Unlike for K^+^, the site is occupied by a single Ca^2+^ ion (Fig. 5h). In the presence of Ca^2+^, membrane thinning near the open vestibule is more frequent in POPC membranes (7 of 10 trajectories) than in DO-mix bilayers (1 of 10) (Fig. 5 Supp. 1e,f), and when it occurs its extent is comparable to that seen in the presence of K^+^ (Fig. 5l). Furthermore, in the Ca^2+^ simulations, membrane thinning correlates with increased hydration of the ND vestibule (Fig. 5i,j) and with the increased dynamic separation of the TM1-TM2 helices (Fig. 5l, Fig 5 Supp. 1h). Finally, membrane thinning is absent in the Ca^2+^ simulations of hXkr4^α^ (Fig. 5k, m, Fig. 5 Supp. 1g). Consistently, in these trajectories the vestibule remains poorly hydrated (Fig. 5i,j), and the TM1-TM2 distance remains short (Fig. 5m). These results suggest that the ND vestibule can bind K^+^ and Ca^2+^, and that both ions exert similar effects on membrane thinning, hydration, and dynamics. The main difference is that Ca^2+^ binding is more stable (Fig. 5 Supp. 5d), and it promotes more frequent membrane bending (Fig. 5 Supp. 5e).

To determine whether Ca^2+^ functionally modulates scrambling by purified hXkr4, we performed the in vitro scrambling assay in 2 mM EGTA to buffer free Ca^2+^ <10 nM [48], with 0.5 mM Ca^2+^, and in unbuffered conditions, where the free Ca^2+^ concentration is ∼10 µM [48]. Our results show that a ∼50,000-fold change in the free Ca^2+^ concentration has no measurable effect on the scrambling rate constants (Fig. 5n, Fig. 5 Supp. 1i), indicating that Ca^2+^ is not a required activator for scrambling by hXkr4, at least in the presence of 300 mM K^+^. We note that although in our simulations Ca^2+^ promotes more frequent membrane thinning than K^+^ and has high occupancy for the ND vestibule cation site, these effects likely reflect the high Ca^2+^ concentrations used in our simulations to enhance the frequency of spontaneous ion binding. Further, in our experiments the K^+^ concentration is ∼600-fold higher than that of Ca^2+^. Thus, the lack of functional modulation by Ca^2+^ likely reflects that during our scrambling assays the ND vestibule cation site is occupied by K^+^ which can also facilitate membrane thinning. Together, these results suggest that the ND vestibule forms a site that can bind both mono- and divalent cations.

## Discussion

The Xkr apoptotic scramblases, the human Xkr4, -8, and -9 and the nematode CED-8 [14, 15], play a key role in the recognition and clearance of apoptotic cells by macrophages. However, the mechanisms underlying their activity and regulation remain poorly understood. The current proposal is that Xkr activation entails cleavage of their N- or C-termini by effector caspases [14, 15], which induces oligomerization [22, 27, 29] and causes a conformational rearrangement that exposes the charged stairway residues in the ND repeat [30] (Fig. 1 Supp1b). However, the activity of Xkr scramblases is also regulated via unknown mechanisms by cellular factors, such as phosphorylation [27], binding of Ca^2+^ and of a peptide from the nuclear protein XRCC4 [22, 29], or by their integration into complexes with bulk lipid transport proteins [32, 35]. Further, Xkr8 and Xkr9 purify as monomers, and neither shows evidence of oligomerization or of conformational changes following caspase processing [30, 31, 36], suggesting their structures represent inactive states.

Here, we show that two purified apoptotic Xkr scramblases, hXkr4 and CED-8, scramble lipids when reconstituted in liposomes. Their activity does not require caspase processing; rather, both full-length proteins are active (Fig. 1) and, in the case of CED-8, a construct mimicking caspase processing does not have increased activity (Fig. 1). Further, our results show that hXkr4 purifies as a monomer (Fig. 1 Supp. 1a), does not form higher order oligomers when reconstituted in proteoliposomes where it is active as a scramblase (Fig. 1g-h), and its activity is not dependent on Ca^2+^ binding (Fig. 5). Thus, neither caspase cleavage nor oligomerization or divalent binding are required for hXkr4 activation. These conclusions contrast with reports indicating that the activation of hXkr4 requires caspase cleavage, dimerization, as well as binding of Ca^2+^ and of the XRCC4 peptide [22, 29]. While we do not have a definitive explanation for this discrepancy, we speculate that in the complex context of a cell hXkr4 could be inhibited by yet unknown partners, either proteins or lipids, that are lost during purification. Their dissociation could be facilitated by caspase cleavage and/or by the binding of Ca^2+^ and of the XRCC4 peptide, rationalizing the results of the cell-based measurements. Nonetheless, our results show that the minimal functional unit of hXkr4 is the full-length, monomeric, protein and that caspase processing or Ca^2+^ binding are not required for its activity.

In our structure of full-length hXkr4 we observe significant rearrangements compared to the conformations of Xkr8 and -9. The two internal repeats, ND and CD, undergo a rotation around the two-fold axis of symmetry of the protein which results in the opening of the large and hydrophobic transmembrane C1 cavity to the bilayer core. In the Xkr8 and Xkr9 structures, this cavity is closed by the interactions of TM2 and TM3 with IH3 (Fig. 3) and plugged by the short C-terminal helix. While the opened C1 cavity is sufficiently wide to harbor lipids, and indeed we observe a lipid-like density in this region, its hydrophobic nature renders it poorly suited to accommodate hydrophilic lipid headgroups, and thus serve as a scrambling pathway. Indeed, our functional experiments and MD simulations show that the C1 cavity does not play a functional role in lipid scrambling by hXkr4 (Fig. 3, 4, and Fig 4. Supp1a).

In our structure, the rotation of the ND and CD repeats breaks the inter-repeat interactions between TM2 and TM3 in the ND and IH3 in the CD (Fig. 3 Supp. 1d-g; Supp. Movie 1). This disengagement allows the ND repeat to rearrange so that the vestibule, formed by TM1, TM2, and TM3 and harbors the negatively charged stairway residues (Fig. 2c-d), opens to the extracellular solution and its electrostatic profile becomes pronouncedly electronegative (Fig. 2f-h). Importantly, the TM2 helix has now space to move as it is directly exposed to the membrane. Indeed, in our MD simulations of cryoEM hXkr4 the TM2 helix is dynamic and samples conformations where it moves away from TM1 as the ND vestibule becomes hydrated and occupied by cations (Fig. 4). These rearrangements open a membrane-exposed fenestration of the ND vestibule and are associated with a pronounced thinning of the membrane in this region (Fig. 4), which might facilitate scrambling. Notably, these dynamics and accompanying membrane thinning are absent in our αFold2 model of hXkr4, as the tight packing of TM2 against the IH3 of the CD repeat prevents movements and are dampened in DO-mixed membranes where scrambling is reduced (Fig. 4l). Finally, in the conformation with a closed C1 cavity and ND repeat (adopted by Xkr8, Xkr9, and hXkr4^α^) the TM2 helix forms extensive inter-repeat interactions with the IH3 and with the C-terminal helix in the CD (Fig. 3 Supp. 1d-f), and these proteins are inactive. Thus, the rearrangement of the ND and CD repeats in the hXkr4 structure enable dynamic rearrangements and hydration of the ND vestibule, which promote membrane thinning. It is likely that additional rearrangements in the ND are needed to enable scrambling, compared to those seen in the cryoEM conformation. Although our MD simulations show membrane thinning, they do not capture full lipid scrambling events, as the IH1-IH2 hairpin and TM2 form a constriction that prevents the full translocation of the lipid headgroups (Fig. 4h). More extensive sampling is likely needed to capture the full extent of the rearrangements needed for scrambling.

Previous work on TMEM16 proteins established that the key structural feature of lipid scramblases is the presence of a hydrophilic groove that locally thins and distorts the membrane [38, 39, 47, 60–62]. Rearrangements of this groove between open and closed conformations modulate the scrambling activity. Our present findings, together with the structures of Xkr8 and - 9, suggest that the Xkr scramblases function according to a similar paradigm, with the ND vestibule serving a role reminiscent of the TMEM16 groove. In the inactive Xkr conformation (Xkr8, -9, and the Alphafold model of Xkr4) the ND vestibule is poorly hydrated, non-dynamic, and the charged stairway residues are buried. This conformation is stabilized by interactions of the C-terminal helix with TM1, TM3 and IH3, which are removed upon caspase processing, facilitating activation. In hXkr4, the C-terminus is longer than in Xkr8 and -9, and the caspase cleavage site is more distal from the membrane (Fig. 1 Supp. 1a), suggesting a different mode of regulation. Indeed, in our structure the C-terminus of Xkr4 does not form tight interactions with the transmembrane region of the protein and is poorly resolved (Fig. 2a), indicating it is dynamic. In the active Xkr4 conformation, the ND and CD repeats have separated, and the TM2 is not constrained in position by the inter-repeat interactions (Supp. Movie 1). This, together with the ensuing increased hydration, allows the ND vestibule to become dynamic and sample conformations where the TM1 and TM2 helices become more separated. This leads to an increased electronegative profile of the region, which promotes membrane thinning. Our data suggests that in Xkr4 the charged residues in the ND vestibule reshape the membrane in its vicinity even though they remain buried within the protein. This is unlike what is seen in the TMEM16s where the open hydrophilic groove is directly exposed to the membrane core [38, 39, 47, 52, 55, 61]. We hypothesize that the electric field created by these charged residues can reshape the membrane, even though these side chains remain buried, as it is less dampened in the low dielectric environment of the bilayer core than it would be in water (Fig. 4). In support of our hypothesis, we note that charge-neutralizing mutations of these residues in hXkr4 reduces the frequency of membrane thinning in our simulations (Fig. 5a-g), and equivalent mutations severely impair scrambling by Xkr8 in cells [30].

In summary, we showed that monomeric, full-length hXkr4 is an active phospholipid scramblase and that its activity is regulated by membrane properties, in a manner reminiscent of TMEM16 scramblases [38, 39, 47]. The reduced inter-repeat interface of the hXkr4 conformation allows the ND vestibule to become hydrated and dynamically rearrange to induce membrane thinning. [38, 39, 47, 60, 62]. While more work is needed to elucidate the precise mechanisms of Xkr4 regulation by caspase processing and ligand binding, our results suggest that the unusual architecture of hXkr4, with several acidic residues buried within the ND repeat, promotes membrane thinning which might facilitate lipid scrambling. Thus, the ability to thin membranes might be a key mechanistic feature shared by structurally unrelated scramblases.

## Methods

### Expression and purification of human Xkr4

Full length human Xk-related protein 4 (hXkr4) was cloned into a modified pBacMam vector with a C-terminal TEV cleavage site followed by FLAG-6xHis tag [44]. Recombinant hXkr4 protein was expressed in HEK-293F suspension cells following baculovirus mediated mammalian cell expression system [44]. 100 ml of P2 generation of viruses were used to infect 1 L of HEK-293F suspension cells (cell density 2.5-3 million/ml) and the cells were kept in a 37 °C incubator shaker for 24 hours with 5% CO_2_ and 110 rpm speed. After 24 hours 10 mM Na-butyrate was added, and the cells were stored in 30 °C incubator shaker for 48 hours with 5% CO2 and 110 rpm speed. After 72 hours of infection, the cell pellet was collected by centrifugation at 2500 rpm.

Cell pellets were resuspended in a lysis buffer containing 300 mM NaCl, 1 mM tris(2-carboxyethyl)phosphine (TCEP), 50 mM HEPES, pH 7.4, protease inhibitor cocktail and trace amounts of DNase. The resuspended cells were sonicated briefly, and the cell debris were discarded by centrifuging at 13000 rpm at 4 °C for 20 minutes. The resulting supernatant was then subjected to high-speed ultracentrifugation at 40000 rpm for 1 hour at 4 °C to isolate membrane fractions. Isolated membrane fractions were homogenized and later solubilized in solubilization/ extraction buffer containing 300 mM NaCl, 1 mM TCEP, 50 mM HEPES, pH 7.4, protease inhibitor cocktail, 2% (w/v) n-Dodecyl-D-Maltoside (DDM) or Lauryl Maltose Neopentyl Glycol (LMNG) and 0.4% (w/v) Cholesteryl HemiSuccinate CHS. The solubilization step was carried out at 4 °C for 2-3 hours with continuous stirring or rotation. The insoluble fractions were discarded by centrifugation at 13000 rpm for 20 minutes. The soluble supernatant was incubated with Flag resin for 2 hours at 4 °C with continuous rotation and were collected on an affinity column by gravity flow. The collected beads were washed with 20 column volumes of wash buffer containing 200 mM NaCl, 1 mM TCEP, 50 mM HEPES, pH7.4, 0.05% DDM-0.01% CHS or 0.001% (w/v)

LMNG-0.0001% (w/v) CHS. The protein was eluted by adding (500 µg/ml) Flag peptide. The eluted fractions were concentrated using a concentrator with MW cut-off 100 kDa and were subjected size exclusion chromatography on a Superose 6 column using SEC buffer containing 150 mM NaCl, 1 mM TCEP, 20 mM HEPES, pH7.4, and 0.05% DDM-0.01% CHS or 0.00075% LMNG-0.000075% CHS. Xkr4 used for native mass spectrometry (nMS) was expressed in GnTI-cells to reduce glycosylation. The protein was purified following as described above but using 0.02% (w/v) DDM-0.004% (w/v) CHS in elution and the following size exclusion chromatography.

### Expression and purification of CED-8

The full length CED-8 from *C. elegans* was cloned into a modified pFastBac vector with a C-terminal TEV cleavage site followed by GFP-FLAG-6xHis tag. Protein was expressed in High Five cells following baculovirus mediated insect cell expression system [63, 64]. 10-20 ml of P2 generation of viruses were used to infect 1 L of Hi5 suspension cells (cell density 1.5-2 million/ml) and the cells were kept in a 27 °C incubator shaker for 72 hours with 110 rpm speed. After 72 hours the cell pellet was collected by centrifugation. Cell pellets were resuspended in a lysis buffer containing 50 mM HEPES pH 7.4, 300 mM NaCl, 1 mM TCEP and protease inhibitor cocktail (Roche) and trace amount of DNase. The resuspended cells were sonicated briefly, and the cell debris were discarded by centrifuging at 13000 rpm at 4 °C for 20 minutes. The resulting supernatant was then subjected to high-speed ultracentrifugation at 40,000 rpm for 1 hour at 4 °C to isolate membrane fractions. Isolated membrane fractions were homogenized and solubilized in a buffer containing 300mM NaCl, 50 mM HEPES, pH 7.4, 1 mM TCEP, protease inhibitor cocktail, 2% DDM, and 0.4% CHS. The solubilization step was carried out in 4 °C for 2-3 hours with continuous stirring or rotation. The insoluble fractions were discarded by centrifugation at 13000 rpm for 20 minutes. The soluble supernatant was incubated with Flag resin for 2 hours at 4 °C with continuous rotation and were collected on an affinity column by gravity flow. The collected beads were then washed with 20 column volumes of wash buffer containing 200 mM NaCl, 50 mM HEPES, pH7.4, 1mM TCEP, 0.05% DDM, 0.01%CHS. The protein was isolated after incubating the Flag resins overnight with 5ml of wash buffer and TEV protease at 4 °C. The resulted fractions were then subjected to a nickel-his affinity column to remove TEV protease. The eluted fractions were concentrated using a concentrator with MW cut-off 50 kDa and were subjected size exclusion chromatography on a superose 6 column using SEC buffer containing 150 mM NaCl, 20 mM HEPES, pH7.4, 1 mM TCEP and 0.05% DDM-0.01% CHS. The ΔCED-8 construct, corresponding to residues 22-420 of CED-8, was expressed and purified following the same protocol.

### Liposome reconstitution

Liposomes were prepared as described [48] using the following lipid compositions: a 7:3 mixture of 1-palmitoyl-2-oleoyl-glycero-3-phosphocholine (POPC, 16:0-18:1) and 1-Palmitoyl-2-oleoyl-sn-glycero-3-[phospho-rac-(1-glycerol)] (POPG 16:0-18:1) (POPC:POPG); a 7:3 mixture of 1,2-dioleoyl-sn-glycero-3-phosphocholine1 (DOPC, 18:1), 2-dioleoyl-sn-glycero-3-phosphoglycerol (DOPG, 18:1); (DOPC:DOPG); a 2:1:1 mixture of 1,2-Dioleoyl-sn-glycero-3-phosphoethanolamine (DOPE 18:1/18:1 PE), DOPC, and 1,2-dioleoyl-sn-glycero-3-phospho-L-serine (DOPS 18:1 PS) (DOPE:DOPC:DOPS); a 2:1:1 mixture of 1-palmitoyl-2-oleoyl-sn-glycero-3-phosphoethanolamine (POPE 16:0-18:1 PE), POPC, and 1-palmitoyl-2-oleoyl-sn-glycero-3-phospho-L-serine (POPS 16:0-18:1 PS) (POPE:POPC:POPG), and soybean polar lipid. Chain length experiments were carried out using a 7:3 PC:PG lipid headgroup background and the following acyl chains [38]: a mixture of 50% 1,2-dimyristoyl-sn-glycero-3-phosphocholine (DMPC, 14:0) and 1,2-dimyristoyl-sn-glycero-3-phospho-(1′-rac-glycerol) (DMPG, 14:0) with 50% POPC:POPG (C14 membrane); DOPC:DOPG (C18 membrane); 1,2-dieicosenoyl-sn-glycero-3-phosphocholine (20:1 PC), 1,2-dieicosenoyl-sn-glycero-3-(1′-rac-glycerol) (20:1 PG) (C20 membrane); 1,2-dierucoyl-sn-glycero-3-phosphocholine (DEPC, 22:1) and 1,2-dierucoyl-phosphatidylglycerol (DEPG, 22:1) (C22 membrane). Lipids were dissolved in chloroform, including 0.4% w/w tail labeled 1,2-dipalmitoyl-sn-glycero-3-phosphoethanolamine-N-(7-nitro- 2-1,3-benzoxadiazol-4-yl) (NBD-PE), were dried under N_2_ gas. The resulting lipid film was washed with pentane, dried under N_2_ gas, and resuspended at 20 mg/ml (for soybean polar 10 mg/ml) in buffer containing 300 mM KCl, 50 mM HEPES pH 7.4 with 35 mM 3-[(3- cholamidopropyl) dimethylammonio]-1-propanesulfonate (CHAPS). The mixture was sonicated until clear. Protein was subsequently added at a concentration of 5 µg protein/mg lipids. Detergent removal was carried out by using Bio-Beads SM-2 (Bio-Rad) with rotation at 4 °C. For all mixtures, except for one containing POPE, 5 exchanges of 200 mg ml^-1^ Bio-Beads were used. For the POPE mixture, 4 exchanges of 150 mg ml^-1^ Bio-Beads were performed. Calcium or EGTA were introduced using sonicate, freeze-thaw cycles. The liposomes were extruded 21 times through a 400-nm membrane before use.

### In vitro scrambling assay

In vitro scrambling assay was performed as described [45]. Liposomes were extruded 21 times through a 400 nm membrane prior to use. 20 µl of liposome were then added to a final volume of 2 mL of buffer containing 300 mM KCl, 50 mM HEPES pH 7.4. The fluorescence intensity of the NBD (excitation-470 nm emission-530 nm) was monitored over time with mixing using a PTI spectrophotometer. After 100 seconds, sodium dithionite was introduced at a final concentration of 40 mM. Data acquisition was done using the FelixGX 4.1.0 software at a sampling rate of 3 Hz.

### Quantification of scrambling assay

Quantification of the scrambling assay and determining the scrambling rate constants were done as described [45]. In brief, the fluorescence decay time course was fit to the following equation

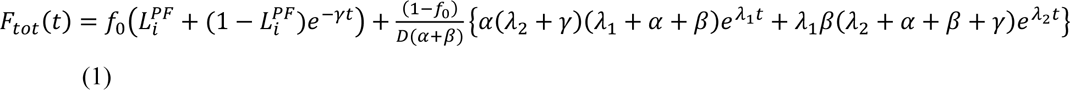

Where

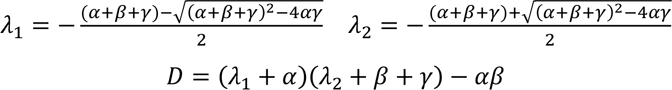

and F_tot_(t) is the total fluorescence at time t, L_i_^PF^ is the fraction of NBD-labeled lipids in the inner leaflet of protein free liposomes, where γ is the rate constant of dithionite reduction, f_0_ is the fraction of protein-free liposomes in the sample, α and β are respectively the forward and backward scrambling rate constants. The free parameters of the fit are f_0_, α and β while L_i_ ^PF^ and γ are experimentally determined from experiments on protein-free liposomes. In protein-free vesicles a very slow fluorescence decay is visible, likely reflecting a slow leakage of dithionite into the vesicles or the spontaneous flipping of the NBD-labeled lipids. A linear fit was used to estimate that the rate of this process is L = (5.4±1.6)10-5 s^-1^ [45]. For Xkr4 functional data in the PM-like condition, f0 was set to free to reflect the low reconstitution efficiency with this lipid composition. In the case of C18 and C22 lipids when the reconstitution efficiency was as high as that in C14, f_0_ from C14 was used for data analysis. Data was analyzed using the custom program Ana (available at http://users.ge.ibf.cnr.it/pusch/) and Prism 7.0 (GraphPad, San Diego, CA) or SigmaPlot 10.0 (SYSTAT Software).

### Sample preparation and optimization

To freeze grids, the monomeric hXkr4 peak fractions were concentrated to 3.5-4 mg/ml immediately after SEC using a concentrator with MW cut-off 100 kDa. Grids were prepared as follows: 3.5 μL of hXkr4 (4 mg/mL) were applied to a glow-discharged Quantifoil (Au 1.2/1.3 200 mesh) grid, incubated for 3 seconds at 100% humidity and 4 °C, blotted for 3 seconds with a blot force -4 and plunge frozen in liquid ethane using a Vitribot Mark IV (FEI). Images were acquired on a 300 kV Titan Krios microscope (Thermo Scientific) equipped with a K3 direct detection camera (Gatan) at NYU Langone Health’s cryo-Electron Microscopy Laboratory.

### Preparation of nMS ready proteoliposomes and downstream nMS experiments

We used our previously developed protocol for preparing nMS-ready proteoliposomes [40, 41]. Briefly purified Xkr4 was reconstituted in PM-like and DOM-mix liposomes using a Sephadex G50 column. The Sephadex G-50 powder was dissolved in ammonium acetate buffer and sonicated in a water bath for 5 min. This suspension was then swelled overnight while being degassed under a vacuum. On the day of the experiment, the Sephadex column was prepared by filling an empty column packed with the pre-swollen Sephadex gel. Separately, dried lipid film was resuspended in ammonium acetate buffer (200 mM ammonium acetate, 2 mM DTT). Then, the solution was sonicated for 15 min in a bath sonicator, and 10 freeze–thaw cycles were performed (liquid nitrogen was used for freezing, and a water bath set at 50 °C was used for thawing). Then the appropriate detergent was added, to a final concentration of 2✕ CMC (critical micelle concentration). This solution was then kept on ice for 30 min. After a 30-minute incubation, the desired amount of protein in 2✕ CMC detergent was added, and the mixture was incubated on ice for 2 h. This sample was placed on the prepared column and separated through gel filtration to collect the proteoliposome fraction. All liposomes were prepared using 1% fluorescent lipid to conveniently track the elution of the liposomes. To achieve stable electrospray ionization, in-house nano-emitter capillaries were used with the Q Exactive UHMR (Thermo Fisher Scientific). These nano-emitter capillaries were created by pulling borosilicate glass capillaries (O.D – 1.2mm, I.D – 0.69mm, length – 10cm, Sutter Instruments) using a Flaming/Brown micropipette puller (Model P-1000, Sutter Instruments). A platinum wire was used for all nMS electrospray. For the nMS of proteins from lipid vesicles, the prepared proteoliposomes were used to fill the nano-emitter capillary, which was installed into the Nanospray Flex ion source (Thermo Fisher Scientific). The MS parameters were optimized for each sample. The spray voltage ranged between 0.9 – 1.2 kV, the resolving power of the MS was in the range between 3K – 6K, the ultrahigh vacuum pressure was in the range of 5.51e^-10^ to 6.68e-10 mbar, and the in-source trapping range was between −50V and −250V. The HCD voltage was optimized for each sample ranging between 0 to 200V. All the mass spectra were visualized and analyzed with the Freestyle (ThermoFisher Scientific) software. UniDec [65] was used for the final mass calculation and assembled into figures using Adobe illustrator.

### Data acquisition and processing

Micrographs were acquired on a Titan Krios microscope (Thermo Scientific) with a K3 direct electron detector (Gatan) at NYU Langone Health’s cryo-Electron Microscopy Laboratory. Images were collected with a total exposure time of 2s, total dose of 58.28 e−/Å2, and a defocus range of 0.5 µm to 2.5 µm. Very stringent data collection and processing criteria were used: only micrographs from regions with ice thinner than 70 nm (majority had 15-40 nm thickness) were imaged and only those with contrast transfer function (CTF) estimates <4.0 Å were used during processing [66]. Motion correction, CTF estimation, automated particle picking, and extraction was carried out in *Warp* [67]. Frames were aligned using Motioncorr2 1.4 under control of Appion [68]. Dose weighting was applied according to the dose calibrated in Leginon [69]. Images were tiled into 7x5 regions for optimal alignment. Global and local B-factors were 500 Å^2^ and 100 Å^2^ respectively. The super-resolution images were Fourier binned by 2 to the physical pixel size of 0.825 Å during the alignment. Image quality was monitored by calculating on-the-fly CTF fitting using CTFFIND4 [66]. Aligned and does weighted images were imported into *Warp* [67]. Particles were picked using an expected particle size of 150 Å and a box size of 320 pixels. Local CTF was estimated in *Warp* [67] by tiling images into 7x5 pieces. Image stacks were directly imported into cryoSPARC v3 [54] for processing.

A total of ∼5.5 million particles selected by *Warp*[67] were imported to cryoSPARC and were subjected to extensive 2D classification (>20 rounds) until clear, distinguishable density for the transmembrane domain was visible. Following the 2D classifications, ab initio reconstructions were conducted for more than 8 times with gradually decreasing the maximum resolution from 12 to 4 Å [54]. The resulting model comprising of 357,599 particles was used for heterogeneous refinement while a non-protein like density, most likely an empty detergent micelle, was used as a decoy class. For heterogeneous refinement, the particles from the second round of selected 2D classes were chosen (total particles 4,973,462). After more than 20 rounds of heterogeneous refinement coupled with several iterations of non-uniform refinements, ∼450k particles were selected for a final non-uniform refinement with low pass filter 6 Å ultimately yielded the final model[70–73]. A final resolution of 3.67 Å was determined using the gold-standard Fourier shell correlation (FSC)= 0.143 criterion using cryoSPARC (Fig. 2 Supp. 1f). This map was of sufficient quality to permit building of an atomic model for the TM region of hXkr4.

### Model building

An initial model was generated by Swiss-Model using human Xkr8 (PDB: 7DCE) as the reference. The generated model was then fit into the cryo-EM density model in UCSF Chimera 1.16. The model was refined against density maps using Phenix 1.20 real space refinements with secondary structure restraints and no NCS constraints. The refinements were done for multiple iterations followed by manual curation in WinCOOT 0.9.8. MolProbity was used to estimate the geometric restrains, clash score, and Ramachandran outliers.

### Data availability

Human Xkr4 model and corresponding maps have been deposited to the Electron Microscopy Data Bank (EMDB) and PDB. The depositions include final map, along with sharpened maps, and FSC curve. The EMDB accession code is EMD-44744 and the PDB one is 9BOJ.

### Molecular dynamics simulation

The simulation systems as listed in Supplementary Table 2 were constructed using either one of two different protein models, which are 1) the cryoEM models of hXkr4 of this work (residues 106−167 and 249−516), referred here as to “EM”, and 3) the hXkr4 model (“αFold”) generated by AlphaFold2 [42] Jupyter notebook in ColabFold ver. 1.5.5 at Google colaboratory [74] (https://colab.research.google.com/github/sokrypton/ColabFold/blob/main/AlphaFold2.ipynb), using full protein sequence of hXkr4 (Uniport ID: Q5GH76). Seven missing residues in EM (I367 and E420−I425) were rebuilt using modeller ver. 10.4 [75]. Another 81 missing residues in the cytoplasmic loop between TM2 and IH1 helices (V168−C248), forming an unstructured coil in αFold, were excluded in our simulations. The N- and C-terminal residues of the whole protein (R106 and N516) and the missing gap (S249 and F167) were capped with NH_3_^+^ and COO^-^ groups, respectively. Truncated sidechains of other residues were rebuilt by psfgen tool in VMD software version 1.9.3[76]. Both EM and αFold simulation systems were built using the residues 106−167 and 249−516. Starting from EM, either D125, D129, or E313 was substituted to Ala in the mutant systems. Default protonation state was used in all other ionizable residues. All simulation systems were constructed using membrane builder tool of the CHARMM-GUI website (http://www.charmm-gui.org/)[77], where the protein as a monomer was embedded in a lipid membrane consisting of either 100% POPC (“POPC”) or 50% DOPE:25% DOPC:25% DOPS mixture (“DO-Mix”), solvated with ∼44,000 water molecules. Either 100 mM of KCl (∼80 K^+^ and ∼80 Cl^-^) or CaCl_2_ (∼80 Ca^2+^ and ∼160 Cl^-^) were added in the solution space. All systems started with all ions >25 Å away from D125, D129, and E313. The Ca^2+^ concentration in the simulation was set to be much higher than its physiological range of the extracellular Ca^2+^ concentration, 1∼3 mM, in order to accelerate Ca^2+^ binding from the solution. Each system contains ∼210,000 atoms in total. The details of the system components in all simulation setups are listed in Supplementary Table 2. The simulation box was set to be orthorhombic with periodic boundaries applied at x-y-z axes and dimensions of 140 Å × 140 Å × 110 Å. CHARMM36 force field[78] was employed for the protein, lipids, K^+^, Cl^-^, and TIP3P water model[79]. Ca^2+^ was treated by the multi-site Ca^2+^ model [80], which better reproduces the binding energy between Ca^2+^ and the carboxyl groups of Asp and Glu, and solvation energy and structure of the coordinated water molecules, than Ca^2+^ model in the conventional CHARMM forcefield. The equilibration and production simulations for 10 replicas from each of the systems #1,3−6 in Supplementary Table 2 were performed with Gromacs package ver. 2022.3[81], and 10 replicas from each of the systems #1, 7−9 with Amber ver. 22 [82]. All replicas were generated by assigning initial velocities at 300 K using different random seed at the beginning of the equilibration step. The position restraints on protein and lipid were gradually released during 50 ns equilibration run, followed by 500 ns production run for each replica with time step of 2 fs with constant pressure of 1 atm and temperature of 300 K. The system coordinates of the production run were recorded every 100 ps, leading to 5000 frames per each production run. All other simulation setup details were taken from our previous work [82]. All replicas in each of the systems #1, 2, 5-9 are divided into two subgroups as listed in the column “Subgroups” in Supplementary Table 2, depending on whether the outer leaflet lipid is bent or remains flat during the production run of each replica. The outer leaflet lipid remains flat in all replicas in the systems #3 and 4. The system #1 is divided into four subgroups with different criteria, as listed in the column “Subgroups2”, depending on the outer leaflet lipid bending and the choice of MD software to generate the trajectories.

### Alignment of the MD trajectories for analysis

The trajectories of all replicas of all systems were aligned with all alpha carbons of hXkr4 using the coordinates at *t* = 0 of the equilibration run as a reference. The alignment was performed using Gromacs tool (gmx trjconv). After alignment with hXkr4, the average z coordinate of all phosphorus atoms in the phosphate groups of all lipid heads in both outer and inner leaflets, which is defined as the z center of the bilayer (*z_center_*), was calculated from the production run trajectories of all replicas of the subgroups with the outer leaflet flat in Supplementary Table 2, then the whole system coordinates in the production run trajectories were shifted in the z direction by –*z_center_* in the later trajectory analysis.

### Trajectory of bending of the outer and inner leaflet lipids

An individual lipid molecule was determined to belong to outer (inner) leaflet if the z coordinate of its phosphorus atom at *t* = 0 of the equilibration run was higher (lower) than the –*z_center_*. The assignments of the outer and inner leaflet for individual lipid molecules were kept fixed throughout the whole production run trajectories, regardless of their positions after *t* = 0. The outer leaflet remained flat in the equilibration runs in all systems. The average and standard deviation of the lowest (highest) z coordinates among all phosphorous atoms was calculated for outer (inner) leaflet lipids. Each replica was assigned into the “bent” subgroup, when the lowest z coordinates among all phosphorous atoms of all outer leaflet lipids remained lower than the average by more than three times of the standard deviation from the average, continuously for longer than 10 ns, otherwise assigned into the “flat” subgroup, as defined in the column “Subgroup” in Supplementary Table 2.

### Binding of cation at the cation binding site

Either K^+^ or Ca^2+^ was determined to be bound at the cation binding site, when the ion was located within 6 Å from carboxyl carbons of either D125, D129, or E313 sidechains. The number of bound cations was calculated every MD frame in all replicas.

### The two-dimensional (2D) distribution of the outer and inner lipid heads on the x-y plane

The average z coordinate of phosphorous atoms of either outer or inner leaflets was calculated at 2 Å × 2 Å square grids spanned in the range of 10 Å < x < 120 Å and 10 Å < y < 120 Å on the x-y plane of the simulation box. The average z coordinate of each grid was scaled by color from red to blue, as the z coordinate changes from z = 3 to 23 Å for the outer leaflet, and z= −23 to −13 Å for the inner leaflet. The color scales in the plots of the outer and inner leaflets were set to change in the opposite direction, so that the color on the grid turns red, as the lipid head in both outer and inner leaflets is bent towards *z_center_*. The plot was made using the grid squares where the number of phosphorus atoms was non-zero in more than 0.4 % of the MD frames of each replica. This threshold value was chosen to exclude grid squares with poor sampling of lipid occupancy due to the rough protein-lipid boundary [83]. In grid squares with lower than 0.4% occupancy the standard deviation of 𝑧 was greater than 3 Å. A one-dimensional (1D) cross-section of 2D plot was obtained from x = 10 Å to 55 Å, while y is fixed at y = 68 Å for the outer leaflet, and y = 70 Å for inner leaflet. The fixed y values in the 1D plots were chosen where the difference of the average z coordinates between x = 10 Å and 55 Å was the greatest over all y.

### Water occupancy profile along the TM1-TM2-IH1 groove

A cylinder was defined at the groove between TM1, TM2, and IH1 helices (See Fig. 4 Supp1a), with radius of 8 Å, where the cylinder axis (*h*) was defined as a vector connecting from the midpoint A of the positions of three alpha carbons of Y137, R142, and I324, located at the extracellular side of TM1, TM2, and TM3 helices respectively (the midpoint A was set to be the origin of the cylinder axis, *h* = 0 Å), to the midpoint B of the positions of three alpha carbons of Y122, G155, L264 of TM1, TM2, and IH1 helices respectively, which are located near a short loop between IH1 and IH2 (*h* = 22.2 Å). Then, the cylinder was extended in both directions between *h* = `-`18 Å and 68 Å, where the cylinder reached the extra- and intracellular solution space, respectively. Then, the cylinder was divided into 2 Å-thick slice along its axis between *h* = `-`18 Å and 68 Å, and the average number of water oxygen atoms within each slice was calculated from the production run trajectories using VMD software ver. 1.9.3[76]. The VMD scripts for analyzing the trajectories of the outer (inner) leaflet lipid and water occupancy in the cylinder are available at dx.doi.org/10.6084/m9.figshare.25892728. The ND vestibule around D125, D129, and E313 was defined as *h* = 16 ∼ 20 Å of the cylinder, as shown in Fig. 4 Supp1a.

### Outer leaflet bending as a function of opening of the TM1-TM2 groove and hydration at the ND vestibule

Three variables, which are 1) the distance between alpha carbons of V126 in TM1 and V152 in TM2 helices, 2) the number of water molecules at the ND vestibule around D125, D129, and E313, and 3) the z coordinate of the lowest outer leaflet head, were calculated at every MD frame. The variables #1 and 2 were set as the horizontal and vertical axes in the 2D plot, which were divided in grids with the size of 0.25 Å × 1 in the range of 4.75 Å < x < 21.0 Å and 0 Å < y < 30 Å. The variable #3 was averaged for each grid and for all replicas of each system. Each grid was colored in scale from red to blue, as the variable #3 changes from z = `-`3 to 12 Å.

## Acknowledgements

The authors thank members of the Accardi lab and Boudker lab for helpful discussions and suggestions. The work was supported by National Institutes of Health (NIH) Grant R01AI178180 (to A.A.), R01GM141192 (to K.G.), P41GM116799 (to Wayne Hendrickson, support for R.B.), and the 1923 Fund (to G.K). Some of this work was performed at the Simons Electron Microscopy Center and National Resource for Automated Molecular Microscopy located at the New York Structural Biology Center, supported by grants from the Simons Foundation (SF349247), NYSTAR, and the NIH National Institute of General Medical Sciences (GM103310). Part of the work was performed at NYU Langone Health’s Cryo-Electron Microscopy Laboratory (RRID: SCR_019202) with the help of Dr. Bing Wang and Dr. William Rice, and at the Cryo-EM Core Facility at Weill Cornell Medical College with the help of Dr. Carl Fluck. The authors are grateful for the computational resources under Projects BIP109 (INCITE) and BIP237 (SummitPlus) at the Oak Ridge Leadership Computing Facility, which is a DOE Office of Science User Facility supported under Contract DE-AC05-00OR22725, and for the in-house computational resources of the David A. Cofrin Center for Biomedical Information in the Institute for Computational Biomedicine and the Scientific Computing Unit at Weill Cornell Medical College.

## Author contributions

S.C., Z.F., S.L., and A.A. designed the experiments; R.B. performed initial expression screening; S.C. expressed and purified proteins, performed initial functional characterization and determined hXkr4 structure; Z.F. performed functional experiments and analyzed the data; S.L. and O.E.A. performed MD simulations; S.L. designed and carried out analysis of MD simulations; G.K. contributed resources; K.G. and A.P. designed and carried out M.S. experiments; A.A. oversaw project and wrote the initial draft of the manuscript. All authors edited the manuscript.

## Data availability

The data that support this study are available from the corresponding author upon request. All models and associated cryoEM maps have been deposited into the Electron Microscopy Data Bank (EMDBID: EMD-44744) and the Protein Data Bank (PDBID: 9BOJ). The depositions include final maps, unsharpened maps, local refined maps, and associated FSC curves. Scripts for analysis of MD trajectories are available at dx.doi.org/10.6084/m9.figshare.25892728.

