## Supplementary Materials for "Structure and function of the human apoptotic scramblase Xkr4"

1 Department of Anesthesiology, Weill Cornell Medical College; 2 Physiology, Biophysics and Systems Biology Graduate Program, Weill Cornell Medical College; 3 Nanobiology Institute, Yale University, West Haven, Connecticut 06516, United States; 4 Department of Cell Biology, Yale University School of Medicine, New Haven, Connecticut 06520, United States; 5 Center on Membrane Protein Production and Analysis (COMPPÅ), New York Structural Biology Center, New York, NY 10027, USA, 6 Department of Physiology and Biophysics, Weill Cornell Medical College; 7 Department of Biochemistry, Weill Cornell Medical College

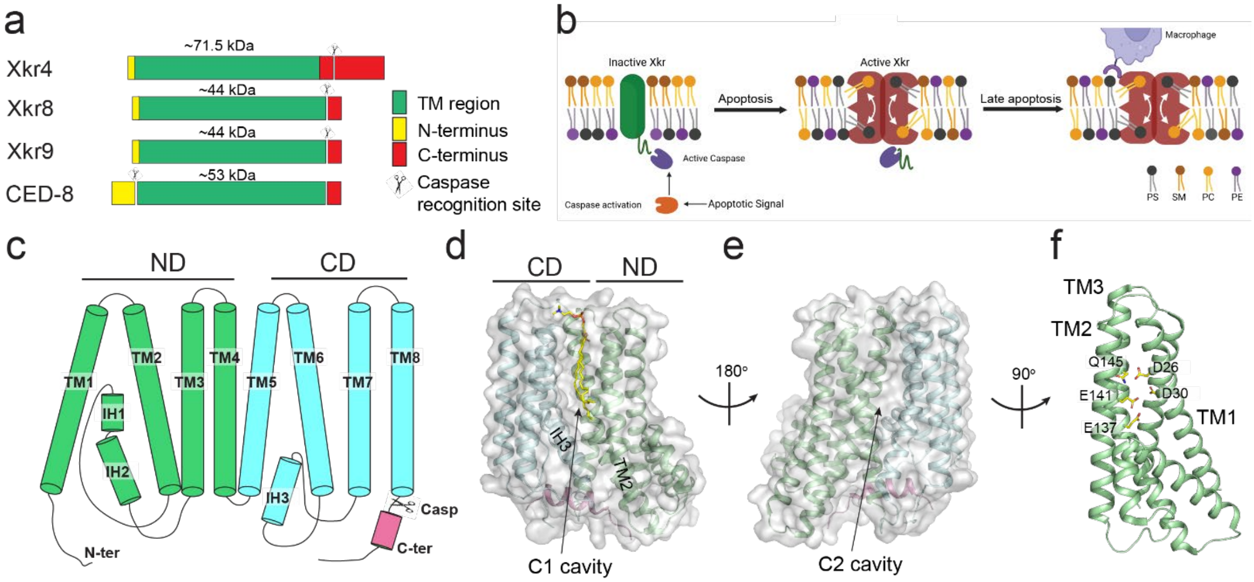


**Figure 1 Supplement 1. Overview of the Xkr4 family.** a) Domain organization of Xkr4, Xkr8, Xkr9 and CED-8. b) Proposed activation mechanism of the apoptotic Xkr scramblases. Initiation of the apoptotic cascade leads to activation of effector caspases (left), which cleave the Xkr C-terminal region to induce dimerization (middle) and a conformational change that allows lipid scrambling (right). c) Schematic topology of Xkr proteins. The ND repeat is in light green, the CD repeat in pale blue and the C-terminal helix in pink. d-e) The structure of hXkr8 (PDBID: 8XEJ) is viewed from the plane of the membrane with the C1 cavity (d) or the C2 cavity (e) in front. The protein is shown in cartoon representation with the ND repeat in light green, the CD repeat in pale blue, the C-terminal helix in pink, and the surface in transparent gray. The lipid in the C1 cavity is in yellow CPK sticks. f) Close-up view of the ND repeat and of the stairway residues, shown in yellow CPK sticks.

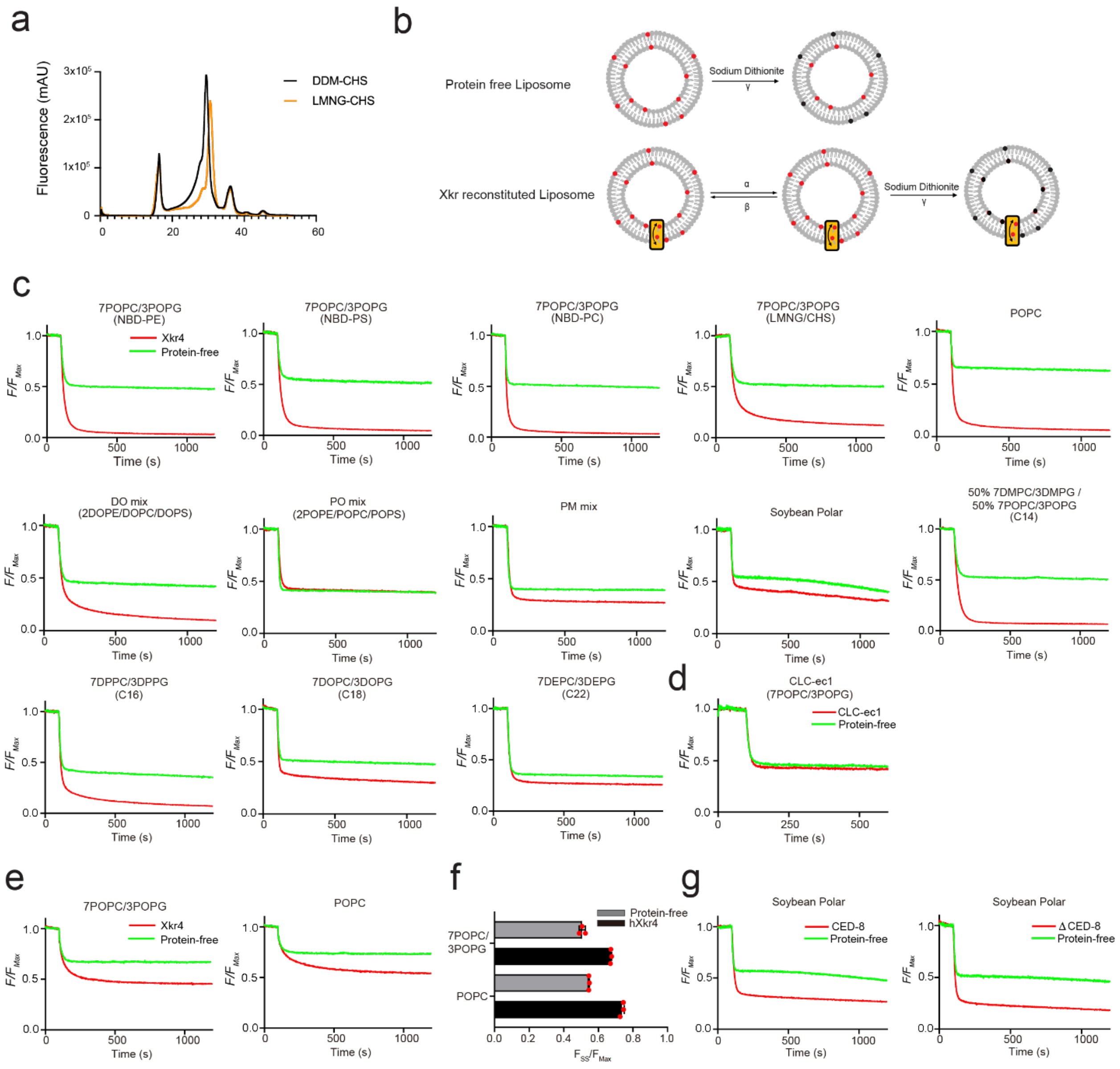

**Figure 1 Supplement 2. Functional characterization of purified hXkr4 and CED-8.** a) Fluorescence size-exclusion chromatography profiles of purified hXkr4 in DDM-CHS (black) and LMNG-CHS (orange). b) Schematic representation of the dithionite-based in vitro scrambling assay. In PF liposomes (top) only outer leaflet NBD-fluorophores (red) are reduced to non-fluorescent (black) by dithionite. In Xkr4-containing proteoliposomes (bottom) NBD-labeled lipids are translocated to the outer leaflet and thus all fluorophores are reduced. c) Representative traces of the dithionite induced fluorescence decay in the scrambling assay for protein free liposomes (green) and hXkr4 (red) reconstituted in vesicles formed from the indicated lipid compositions. d) Representative traces of the dithionite induced fluorescence decay in the scrambling assay for protein free liposomes (green) and CLC-ec1 (red) reconstituted in vesicles formed from the indicated lipid compositions. e) Representative traces of the BSA induced fluorescence decay in the scrambling assay for protein free liposomes (green) and hXkr4 (red) reconstituted in vesicles formed from 7 POPC: 3 POPG (left) or 100% POPC (right). f) Normalized steady state of the fluorescence decay in the BSA back extraction assay in protein free liposomes (gray bar) or hXkr4 proteoliposomes (black bar). Red circles indicate individual experimental values. Data is Mean±St.Dev. g) Representative traces of the dithionite induced fluorescence decay in the scrambling assay for protein free liposomes (green), CED-8 (red), or ΔCED-8 (black) reconstituted in Soybean liposomes.

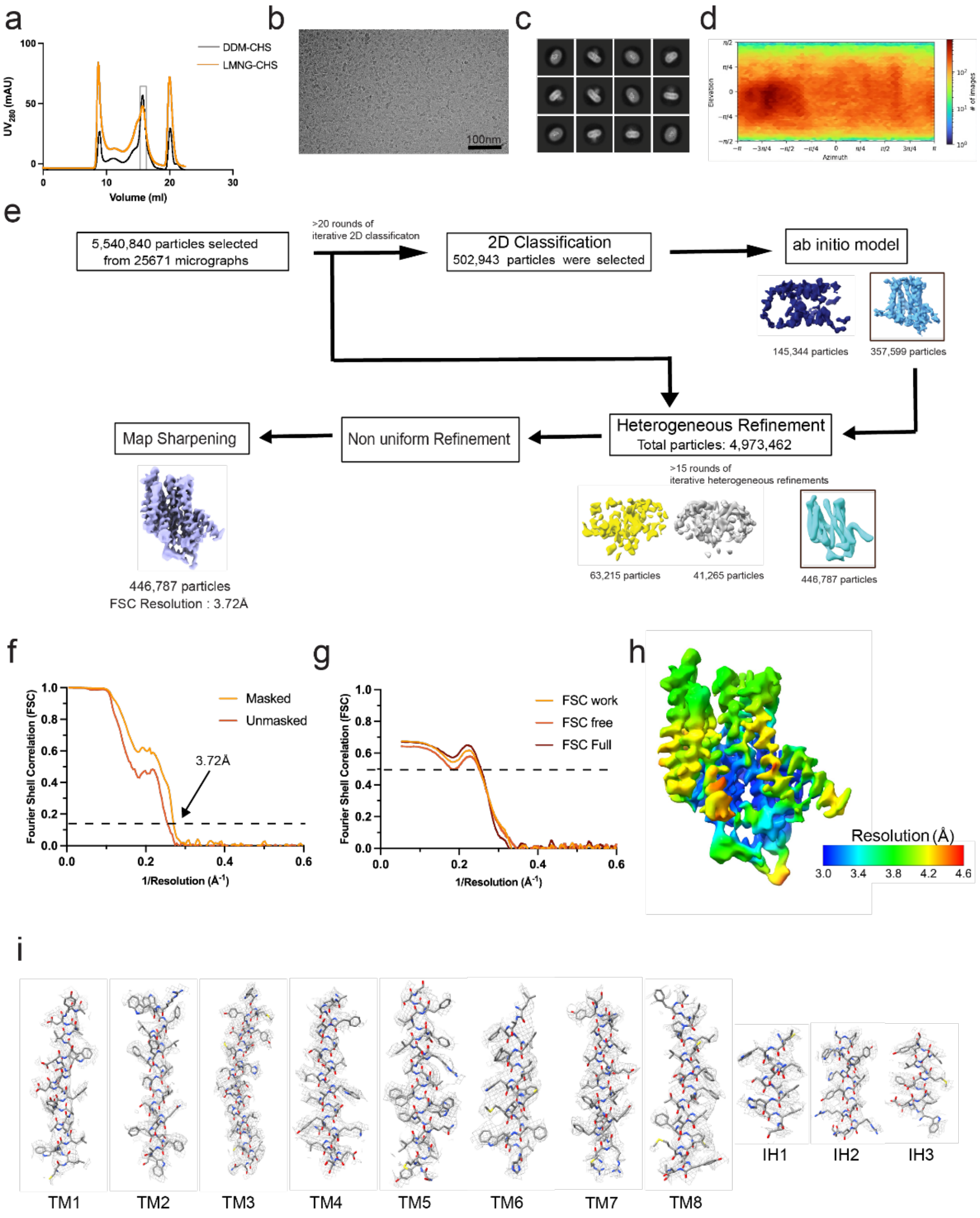


**Figure 2 Supplement 1. CryoEM** image processing scheme for hXkr4. a) Size exclusion chromatography profile of hXkr4 purified in DDM-CHS (black) or LMNG-CHS (orange). Gray box denotes the fractions collected from the LMNG-CHS purification and used for cryoEM imaging. b) Representative cryoEM micrograph of hXk4. c) Representative 2D classes from CryoSPARC. d) Angular distribution of the final reconstruction. e) Summary of the image processing procedure. Classes with black boxes were selected for further processing. f) Fourier shell correlation (FSC) curves for the masked (tight mask from CryoSPARC) and unmasked density maps. g) FSC curves from model–map cross-validation. h) Density map colored by local resolution estimated in CryoSPARC. i) Density maps (mesh) for TM1-10 and IH1-3. Data acquisition and refinement parameters are reported in Table 1.

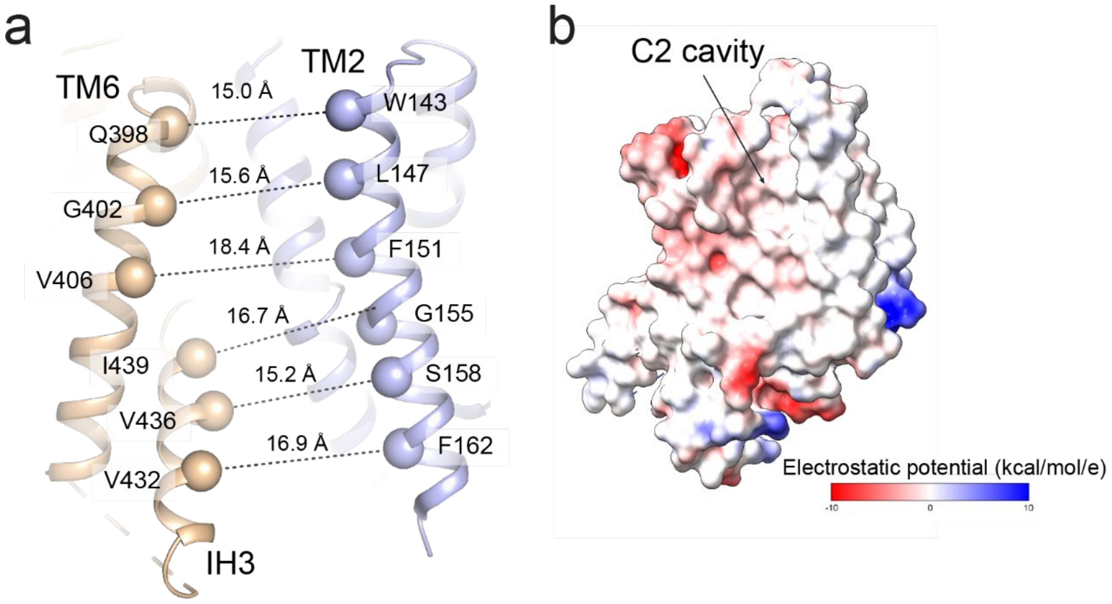


**Figure 2 Supplement 2. The C1 and C2 cavities in hXkr4**. a) Distances of the Cα atoms of residues on TM2 and TM6, IH3 that define the opening of the C1 cavity. b) Electrostatic profile of the hXk4 monomer viewed from the plane of the membrane with the C2 cavity in front.

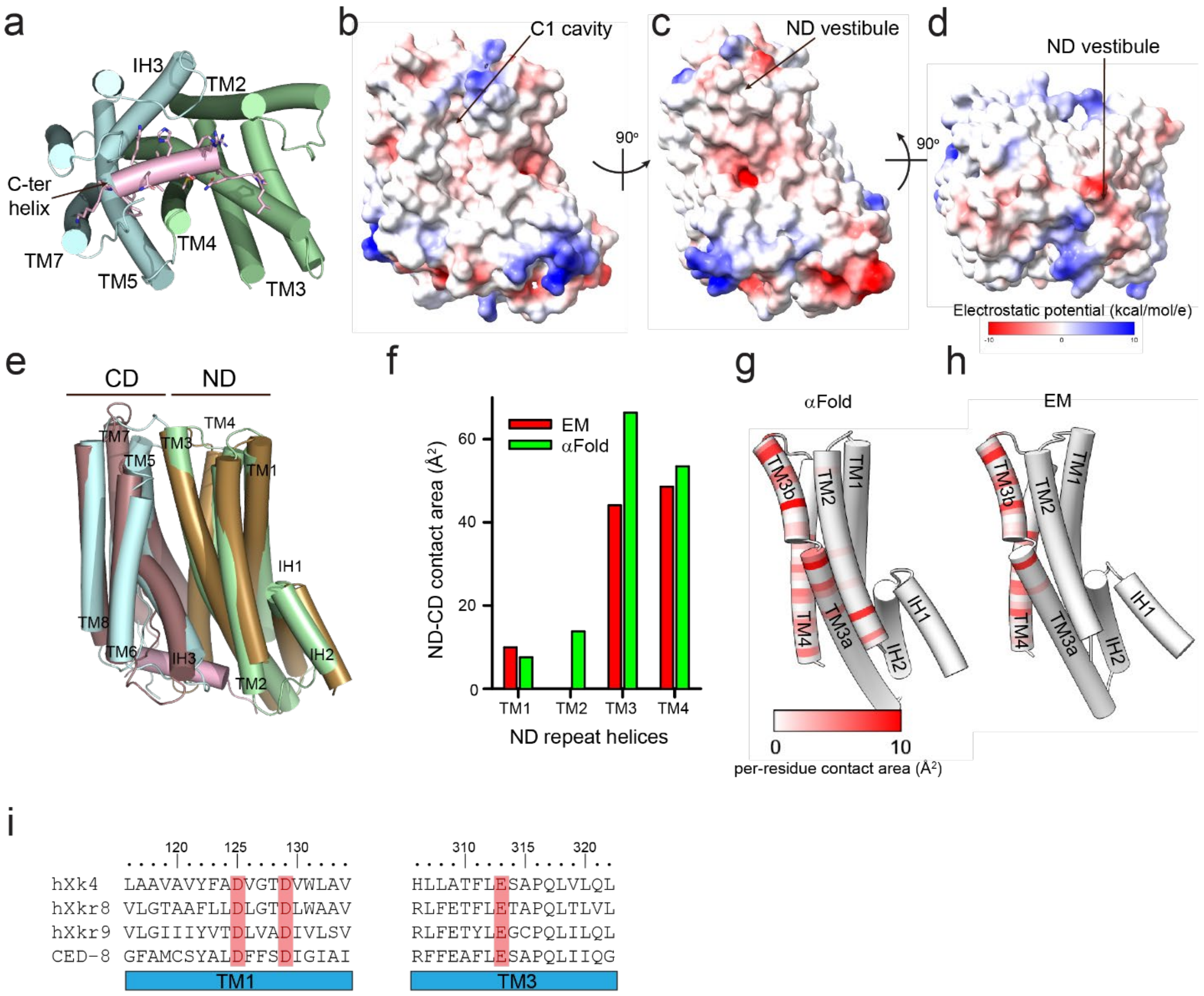


**Figure 3 Supplement 1. Characteristics of hXkr8**. a) The hXkr8 structure in cartoon representation is viewed from the intracellular side. The ND repeat is in pale green, the CD repeat in cyan and the C-terminal helix in pink. Side chains in the C-terminal helix are shown in stick representation. b-d) Electrostatic profile of the hXk8 monomer viewed from the plane of the membrane with the C1 cavity in front (b), from the ND vestibule (c), or from the extracellular side (d) . e) Structural superposition of hXkr8 (PDBID: 8XEJ, ND) and the AlphaFold2 generated model of hXkr4, with the ND colored in green cyan (hXkr8) or maroon (Xkr4α), and the CD in pink f) Per helix surface contact area (in Å$^2$) between the CD and the ND repeat helices (TM1-4), calculated for hXkr4 in the cryoEM (EM, red bars) and AlphaFold2-generated (αFold, green bars) conformations g-h) Per-residue breakdown of the contact surface area in the EM (g) and αFold (h) conformations of hXkr4. i) Conservation of the ND repeat charged residues in hXkr4, hXkr8, hXkr9, and CED-8.

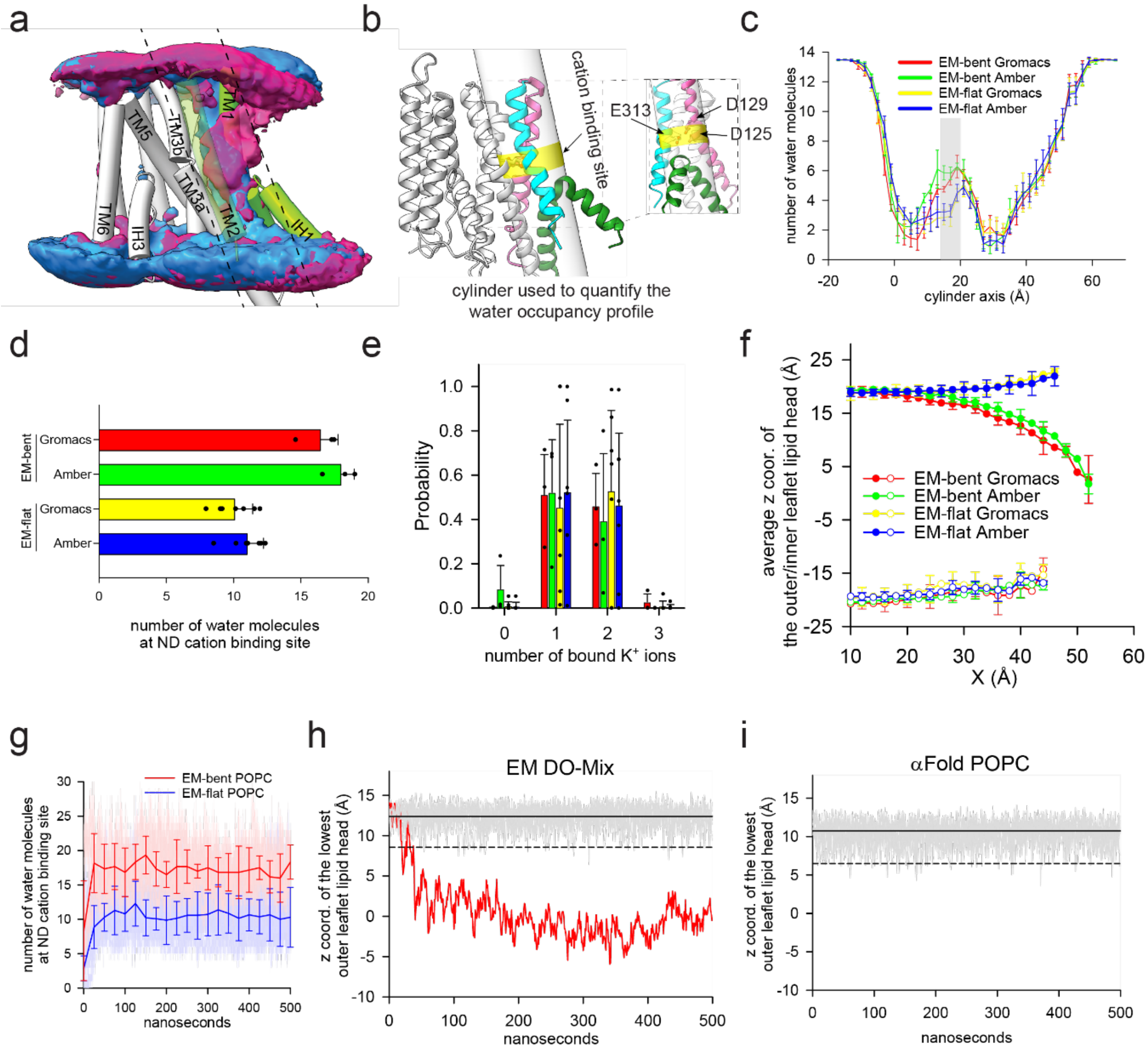


**Figure 4. Supplement 1.** a) The average water density around hXkr4 in MD trajectories of EM-bent POPC (blue) and EM-flat POPC (red), using a threshold of 0.2. The black dashed lines indicate the outline of the cylinder used to quantify water occupancy (see (b)). TM1-2 and IH1-2 of hXkr4 are colored in green, and other helices in light gray. TM2 is set be transparent to show the water density between TM1 and 2. b) The cylinder used to quantify hydration of the ND vestibule. hXkr4 is shown in cartoon with TM1 in pink, TM2 in cyan, IH1-2 in green, and TM3-8 in light gray. The ND vestibule region around D125, D129, and E313 is colored in yellow to indicate the cation binding site. Close-up view of the ND vestibule cation binding site with the sidechains of D125, D129, and E313 shown in stick. c) The average number of water molecules along the cylindrical axis along the ND vestibule for trajectories in POPC of hXkr4 generated using Amber or Gromacs. EM-bent Gromacs (red, n=3), EM-bent Amber (green, n=3), EM-flat Gromacs (yellow, n=7), EM-flat Amber (blue, n=7). The region of the cation site in the ND vestibule (near D125, D129, and E313, 16 Å<h<20 Å) is colored in gray. d-f) Average number of water molecules in the ND vestibule cation site (d), probability distribution of the number of bound $K^+$ ions (e), and average z coordinate of the phosphorous atom of the lowest/highest lipid headgroup from the outer/inner leaflet in MD trajectories of EM-bent Gromacs (red, n=3), EM-bent Amber (green, n=3), EM-flat Gromacs (yellow, n=7), EM-flat Amber (blue, n=7) (f). Data is Mean± St.Dev. g) Time course of the average hydration of the ND vestibule cation binding site in MD trajectories of hXkr4 in POPC membranes of EM-bent (solid red lines) or EM-flat (solid blue lines). Number of water molecules in the ND vestibule of individual replicas are shown in thin solid lines. h-i) Time evolution of the z coordinate of the phosphorous atom in the lowest outer leaflet lipid headgroup of individual trajectories of EM-flat DO-Mix (gray) and EM-bent DO-Mix (red) (h) or of hXkr4$^{\alpha}$ in POPC membranes. Solid lines represent the Avg. and dashed line represent the Avg. – 3xSt.Dev 3 x St.Dev of the z coordinate.

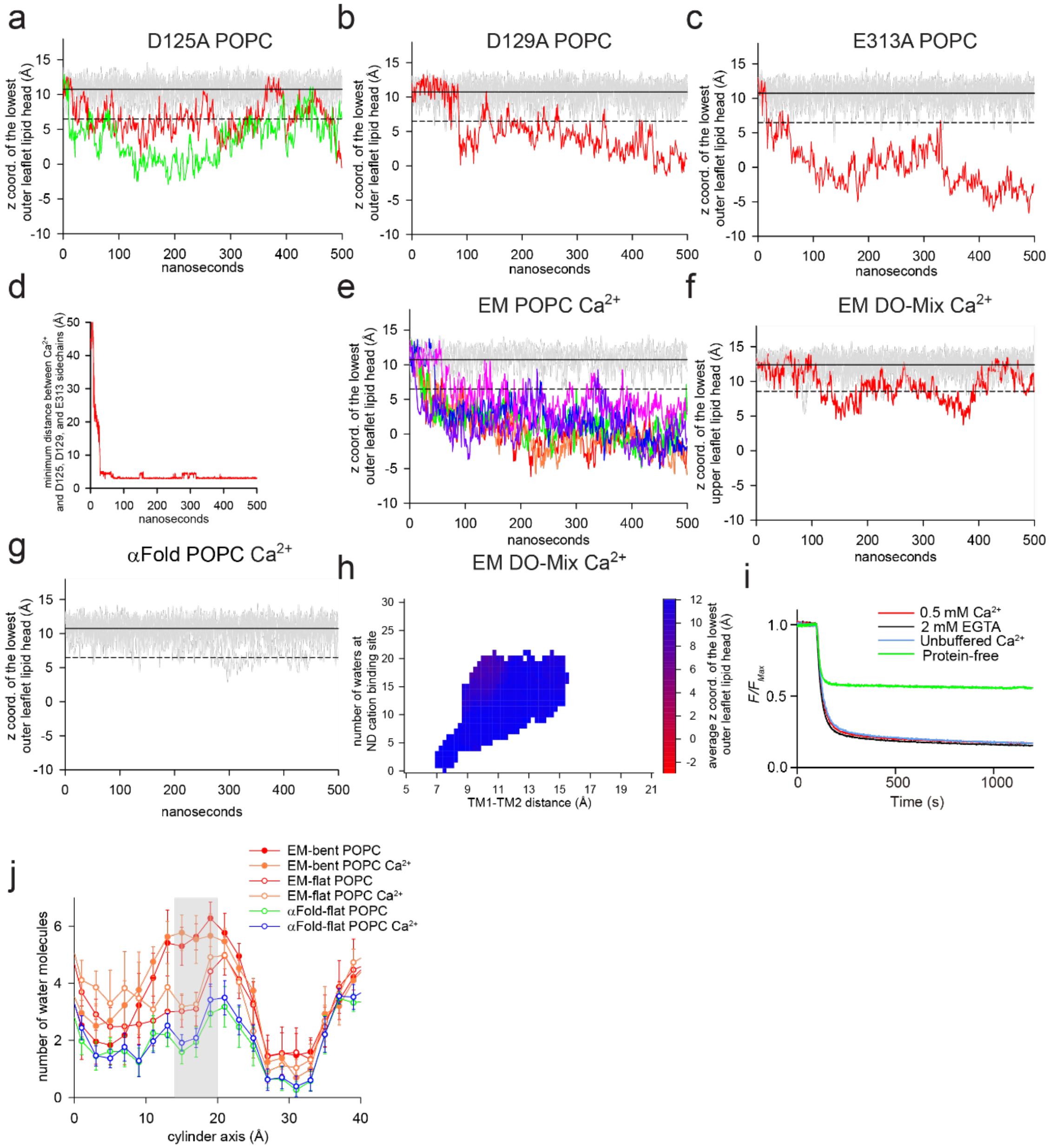


**Figure 5. Supplement 1.** a-c) Time evolution of the z coordinate of the phosphorous atom in the lowest outer leaflet lipid headgroup of individual trajectories of hXk4 D125A (a), D129A (b), and E313A (c) in POPC membranes. Traces where the membrane remains flat are in gray, traces where the membrane bends are colored. Solid and dashed lines represent the Avg. and Avg. – 3xSt.Dev of the z coordinate. (d) Representative trajectory of the minimum distance of a $Ca^{2+}$ ion in the ND vestibule site from the carboxyl carbons of D125, D129, and E313. (e-g) As in (a-c) but for cryoEM hXkr4 (e) and for hXkr4$^{\alpha}$ (f) in POPC membranes in the presence of $Ca^{2+}$, or for cryoEM hXkr4 in DO-Mix membranes in the presence of $Ca^{2+}$ (g). h) The average z coordinate of the phosphorous atom of the headgroups from the lowest outer leaflet lipid is plotted as a function of the V126-V152 Cα distance (x axis) and of the number of water molecules in the cation site in the ND vestibule (y axis) for trajectories for cryoEM hXkr4 in DO-Mix membranes in the presence of $Ca^{2+}$. i) Representative traces of the dithionite induced fluorescence decay in the scrambling assay for POPC protein free liposomes (green), and proteoliposomes reconstituted with hXkr4 in unbuffered $Ca^{2+}$ (blue, ~10 μM), 2 mM EGTA (black, <10 nM), or 0.5 mM $Ca^{2+}$ (red). j) Average number of water molecules in the ND vestibule cation site in MD trajectories of EM-bent (red filled circles) or EM-flat (red empty circles) POPC $K^{+}$, EM-bent (orange filled circles) or EM-flat (red empty circles) POPC $Ca^{2+}$, hXkr4α POPC with $K^{+}$ (green empty circles) or $Ca^{2+}$ (blue empty circles). Shaded area indicates the ND vestibule cation site.

| Lipid | Exact mass (Da) | PM | DO mix |
|---|---|---|---|
| PC(18:1(9Z)/18:1(9Z)) | 785.60 | 23 | 24 |
| PE(18:1(9Z)/18:1(9Z)) | 743.54 | 11 | 50 |
| Cholesterol | 386.35 | 34 | 0 |
| PI(16:0/18:1(9Z)) | 836.52 | 4 | 0 |
| DAG(16:0/18:1(9Z)) | 594.52 | 0 | 0 |
| PS(16:0(9Z)/18:1(9Z)) | 760.52 | 8 | 0 |
| PS(18:1(9Z)/18:1(9Z)) | 787.60 | 0 | 25 |
| SM(d18:1/18:0) | 730.59 | 17 | 0 |
| Brain PIP2 (mixed fatty acyl chains) | 1096.38 (Formula Weight) | 2 | 0 |
| CL(1'-[18:1(9Z)/18:1(9Z)],3'-[18:1(9Z)/18:1(9Z)]) | 1457.04 | 0 | 0 |
| PG(16:0/18:1(9Z)) | 748.50 | | 0 |
| Galactosyl(α) Ceramide (d18:1/16:0) | 699.56 | 0 | 0 |
| Brain Ceramide | 565.95 (Formula Weight) | 0 | 0 |
| Lissamine Rhodamine PE(18:1(9Z)/18:1(9Z)) | 1284.5 | 1 | 1 |

**Supplementary Table 1. Molar percentage of different lipids in PM-like and 2:1:1 DOPE:DOPC:DOPS (DO-mix) liposomes**

| System# | Protein | Cation | Lipid | # of replicas | Subgroups (# of replicas per group) | Subgroups for system#1 (# of replicas per group) |
|---|---|---|---|---|---|---|
| 1 | EM | $K^+$ | POPC | 20 | EM-bent POPC (6)<br>EM-flat POPC (14) | EM-bent Gromacs (3)<br>EM-flat Gromacs (7)<br>EM-bent Amber (3)<br>EM-flat Amber (7) |
| 2 | EM | $Ca^{2+}$ | POPC | 10 | EM-bent POPC $Ca^{2+}$ (6)<br>EM-flat POPC $Ca^{2+}$ (4) | |
| 3 | αFold | $K^+$ | POPC | 10 | αFold -flat POPC (10) | |
| 4 | αFold | $Ca^{2+}$ | POPC | 10 | αFold -flat POPC $Ca^{2+}$ (10) | |
| 5 | EM | $K^+$ | DO-Mix | 10 | EM-bent DO-Mix (1)<br>EM-flat DO-Mix (9) | |
| 6 | EM | $Ca^{2+}$ | DO-Mix | 10 | EM-bent DO-Mix $Ca^{2+}$ (1)<br>EM-flat DO-Mix $Ca^{2+}$ (9) | |
| 7 | EM-D125A | $K^+$ | POPC | 10 | D125A bent (2)<br>D125A flat (8) | |
| 8 | EM-D129A | $K^+$ | POPC | 10 | D129A bent (1)<br>D129A flat (9) | |
| 9 | EM-E313A | $K^+$ | POPC | 10 | E313A bent (1)<br>E313A flat (9) | |

**Table 2. Simulation system setups.** EM, αFold, EM-D125A, EM-D129A, and EM-E313A in the Protein column represent the initial coordinates of each system, which are our cryoEM and alphafold models of hXkr4, and each of alanine mutants generated using EM as a reference model. $K^+$ and $Ca^{2+}$ in the Cation column indicate either 100 mM of KCl or $CaCl_2$ in the solution. The membrane consists of either 100% POPC (denoted as "POPC" in the "Lipid" column) or 50% DOPE:25% DOPC:25% DOPS (denoted as "DO-mix"). Twenty replicas of 500 ns trajectories were generated for system #1 and ten replicas for systems #2-11 using either Gromacs or Amber simulation packages. Each system is divided into two subgroups, bent or flat, depending on whether the outer leaflet is bent in the production run. The subgroup names and the number of replicas assigned to each subgroup are shown in the "Subgroups" column. For system #1, replicas are divided into four subgroups in the "Subgroups2" column, depending on the outer leaflet bending and choice of MD software to generate the trajectory.

**Supplementary Video 1 Morph movie of hXkr4 transitioning from αFold generated model with a closed C1 cavity to the cryoEM conformation with an open C1 cavity.** The protein is viewed first from the plane of the membrane and then from the intracellular side.

**Supplementary Video 2 Morph movie of ND repeat of hXkr4 transitioning from αFold generated model to the cryoEM conformation.** The protein is viewed first from the plane of the membrane.
